# Machine learning-optimized targeted detection of alternative splicing

**DOI:** 10.1101/2024.09.20.614162

**Authors:** Kevin Yang, Nathaniel Islas, San Jewell, Anupama Jha, Caleb M. Radens, Jeffrey A. Pleiss, Kristen W. Lynch, Yoseph Barash, Peter S. Choi

## Abstract

RNA-sequencing (RNA-seq) is widely adopted for transcriptome analysis but has inherent biases which hinder the comprehensive detection and quantification of alternative splicing. To address this, we present an efficient targeted RNA-seq method that greatly enriches for splicing-informative junction-spanning reads. Local Splicing Variation sequencing (LSV-seq) utilizes multiplexed reverse transcription from highly scalable pools of primers anchored near splicing events of interest. Primers are designed using Optimal Prime, a novel machine learning algorithm trained on the performance of thousands of primer sequences. In experimental benchmarks, LSV-seq achieves high on-target capture rates and concordance with RNA-seq, while requiring significantly lower sequencing depth. Leveraging deep learning splicing code predictions, we used LSV-seq to target events with low coverage in GTEx RNA-seq data and newly discover hundreds of tissue-specific splicing events. Our results demonstrate the ability of LSV-seq to quantify splicing of events of interest at high-throughput and with exceptional sensitivity.

## INTRODUCTION

Alternative splicing (AS) varies the exonic and intronic segments included in mature mRNA to substantially increase the diversity of transcript isoforms. Patterns of AS are highly tissue-specific and frequently altered in disease, as revealed by large-scale studies such as the Genotype-Tissue Expression (GTEx) project^1^ and The Cancer Genome Atlas (TCGA)^2^. Mapping the complex landscape of AS has been a major ongoing challenge, driven by key advances in technology.

Short-read RNA sequencing (RNA-seq) is the current most popular approach for transcriptome-wide profiling of AS changes across tissues or conditions. Detection of AS relies on sequencing reads spanning splice junctions, as these provide direct evidence of a particular splicing outcome. However, the majority of RNA-seq reads do not span splice junctions and are thus less informative for AS analysis. To compensate for this bias, robust splicing quantification using RNA-seq typically requires deep sequencing at higher cost compared to gene expression analysis in order to obtain sufficient coverage of splice junctions. Even at near-saturating coverage, RNA-seq still fails to recover certain splicing events which are impeded by issues such as low abundance or secondary structure^3–5^.

Targeted RNA-seq methods provide an efficient way to focus on specific transcripts or RNA regions which cannot be easily studied with standard RNA-seq. A variety of such approaches have been developed, each with different strategies for enriching RNAs of interest. For example, CaptureSeq and TEQUILA-Seq use oligonucleotide probes tiling exonic regions to capture targeted RNAs for sequencing^6,7^. RASL-Seq and TempO-seq use pairs of detector oligos that anneal adjacent to each other on target RNAs and when ligated together, serve as readouts of target RNA abundance^8,9^. Multiplexed PCR is widely used to selectively amplify a set of target cDNAs for either short-read or long-read sequencing^10^. As a promising alternative strategy, Multiplexed Primer Extension sequencing (MPE-seq) is a targeted RNA-seq method for detection of splicing in yeast^11^. MPE-seq performs targeting at the reverse transcription (RT) step, using pools of primers annealing downstream of splice junctions of interest. However, because the complexity of splicing in higher eukaryotes is orders of magnitude greater than in yeast, the use of multiplexed RT for studies of human splicing is significantly more challenging.

Here, we describe Local Splicing Variation sequencing (LSV-seq), a targeted sequencing method we developed to address the limitations of RNA-seq and better capture AS events of interest. Building on prior targeted methods, we designed LSV-seq to offer a unique set of advantages. LSV-seq minimizes the number of required primers per targeted splicing event, enables the discovery and quantification of rare junctions, and precisely discriminates quantitative AS differences in challenging targets. As in MPE-seq, LSV-seq uses customized primer pools to perform highly multiplexed RT adjacent to exon junctions, thereby directly enriching for junction-spanning reads. However, LSV-seq involves several key advances to convert the original multiplexed primer targeting schema into a robust, generalizable methodology. Primers designed with available tools performed inadequately, which led us to create Optimal Prime, a novel machine learning-based primer design algorithm vastly increasing the per-primer targeting efficiency. We also established a webtool based on Optimal Prime for other researchers to design LSV-seq primers. Separately, we created a novel, optimized library preparation protocol and updated our MAJIQ splicing algorithm^12^ for use with LSV-seq sequencing data. To showcase its final capabilities, we benchmarked LSV-seq directly against conventional RNA-seq and leveraged deep learning splicing code predictions to reassess splicing events with low coverage in human GTEx RNA-seq data. Importantly, LSV-seq recovered hundreds of previously unquantified tissue-specific AS variations which were missed due to poor coverage in RNA-seq. We demonstrate that LSV-seq offers an accurate, sensitive and cost-effective method for the study of AS with the ability to target thousands of AS events.

## RESULTS

### Overview of LSV-seq

In order to better capture AS variations of interest, we developed Local Splicing Variation sequencing (LSV-seq) (Fig. 1a). While standard RNA-seq aims to capture all transcripts in an unbiased manner using random N-mer or oligo-dT reverse transcription (RT) primers, LSV-seq instead captures specific RNA regions of interest using complex pools of targeted RT primers. Importantly, LSV-seq is highly scalable and the number of RNA targets can range from hundreds to thousands. For detection of alternative splicing, the designed primers bind to specific target regions adjacent to selected splice junctions of interest (Fig. 1a). The primer design and targeting of splice junctions takes advantage of the Local Splicing Variation (LSV) formulation and detection in MAJIQ^12^ (Fig. 1b). LSVs are defined as all of the splice junctions entering (*i.e.* single target) or exiting (*i.e.* single source) a specific reference exon, thus allowing splicing studies to capture variations with any number of junctions. Under this framework, LSV-seq can capture target LSVs in MAJIQ’s splice graphs, consisting of all of the splice junctions entering the 3’ node of an AS event. Importantly, the target LSV formulation allows the detection and quantification of unannotated and complex splicing variations involving more than two splice junctions using a single primer. While junctions can be targeted individually, our analysis indicates we are able to capture 79% of splice junctions across GTEx tissues using the target LSV formulation (Fig. 1b).

**Fig. 1.**
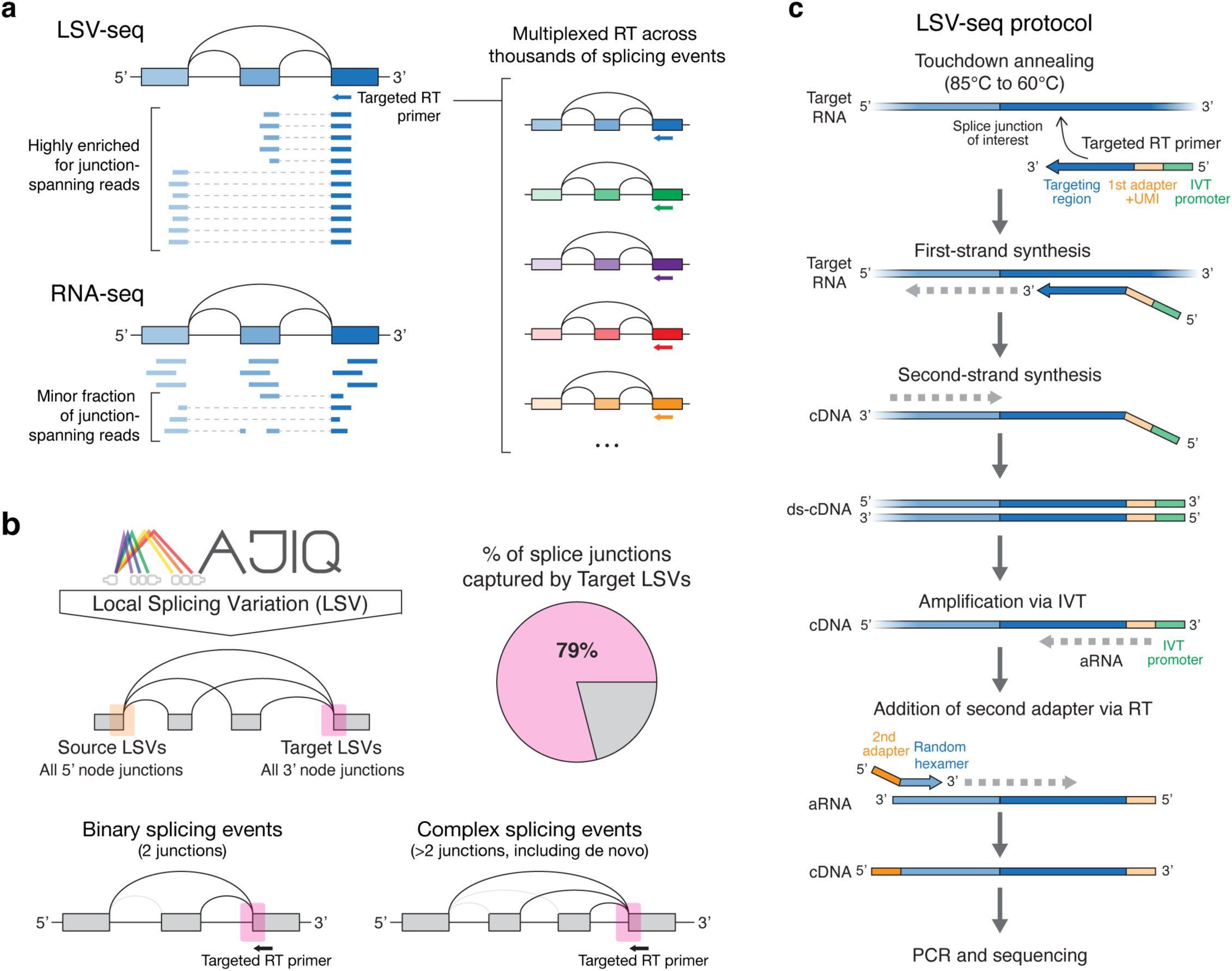
Overview of LSV-seq for targeted detection of alternative splicing. **a**, LSV-seq enriches for junction spanning reads by performing highly multiplexed reverse transcription (RT) with primers anchored directly adjacent to target LSVs. In contrast, only a minor fraction of conventional RNA-seq reads span splice junctions and are informative for splicing quantification. **b**, LSV formulation for splicing events used by the MAJIQ algorithm. Pie chart depicts the percentage of all splice junctions that can be captured by target LSVs. LSV-seq primers can capture both classical binary splicing events as well as more complex events consisting of annotated and novel junctions. **c,** Overview of LSV-seq protocol steps. LSV-seq targeting primers are first annealed to the target RNA using a touchdown protocol to maximize specificity. After first and second strand cDNA synthesis, linear amplification occurs via in vitro transcription (IVT). The amplified RNA (aRNA) then undergoes another RT step to append the second adapter. The resulting cDNA is then PCR amplified and sequenced.

We initially attempted to directly transfer over the previously published MPE-seq protocol^11,13^ from yeast to human cells. However, pilot experiments across two different primer pools resulted in libraries with a low mean percentage of on-target reads ranging from 1.24% to 2.02%, despite substantial amounts of input RNA (50 ug) (Supplementary Fig. S1). We hypothesized that these results were due to the massively increased complexity of the human transcriptome compared to yeast and that further development of the method was needed. Thus, we undertook a series of iterative experimental and computational optimizations focused on improving the specificity and overall yield of the assay, which led to the LSV-seq method presented here.

In our optimized LSV-seq protocol, the targeting primer consists of a constant 5’ sequence which includes a T7 promoter sequence followed by an adapter sequence for PCR amplification and a 10-nucleotide unique molecular identifier (UMI) (Fig. 1c). This invariant region is then followed by the target-specific priming sequence. The LSV-seq protocol then proceeds as follows (Fig. 1c): first, the LSV-seq primer pool is gradually annealed to input RNA using a touchdown step and first-strand cDNA synthesis occurs at an elevated temperature of 60°C to maximize the specificity of the RT reaction. Next, second strand synthesis is performed and the double-stranded DNA then serves as a template for linear amplification by in vitro transcription (IVT). As the targeted RT performed in LSV-seq significantly reduces the amount of starting cDNA compared to conventional RNA-seq, we adopted the IVT step used in single-cell protocols like CEL-seq2^14^ and found that it is critical for increasing the amount of material available for downstream steps. The IVT amplified RNA (aRNA) is then fragmented and proceeds through an additional RT step to append a second adapter sequence. The cDNA is finally amplified by PCR and the resulting library is ready to be sequenced. This final experimental protocol was coupled with optimization of primer design, which we describe next.

### Optimal Prime machine learning models predict high performance primers

We originally attempted to design LSV-seq primers using multiple existing probe design tools^15–17^, but were not able to achieve satisfactory performance across multiple iterations. To address this, we systematically analyzed trends which might reasonably correlate with primer performance. We first compiled an exhaustive dataset of ∼15,000 distinct data points from previously sequenced iterations of LSV-seq libraries. This dataset spanned 5 different primer pools reflecting a logarithmic regime of potential pool sizes, and 5 unique cell line or tissue conditions (Supplementary Table 1). We then defined optimal primer performance in terms of a combined specificity metric and a yield metric, based on rational models of primer binding (Fig. 2a). The first component of the specificity metric is the Fraction of Primer Binding (FPB). FPB is computed by inferring the primer of origin from the 5’ end of the read, allowing us to identify reads with nonspecific partial primer binding during the reverse transcription reaction. The second component of the specificity metric is On-Target Fraction (OTF), which is computed by calculating the ratio of reads mapping to the intended on-target loci compared to undesired off-target loci. OTF and FPB are averaged to create a single combined specificity metric used in downstream analyses. We also defined a yield metric, the log total amplification, as the logarithmic sum of the on- and off-target reads combined. The poor observed correlation (R = 0.23) between the specificity and yield metrics prompted us to develop independent explanatory models for each (Fig. 2b). Validating our initial challenges with using previous probe design tools, commonly used heuristics such as melting temperature, primer length, and off-target alignment scores failed to identify any clear explanatory factors (Fig. 2c). Based on calculated R2 scores, the strongest correlated individual metrics we examined corresponded to a maximal explained variance in performance of only 7.3% for the specificity metric (GC content, R = −0.2707), and 22.4% for the yield metric (melting temperature, R = −0.4729). Moreover, many of the individual factors we examined are highly intercorrelated (for instance GC content and melting temperature), implying that basic regression models would not markedly improve performance.

**Fig. 2.**
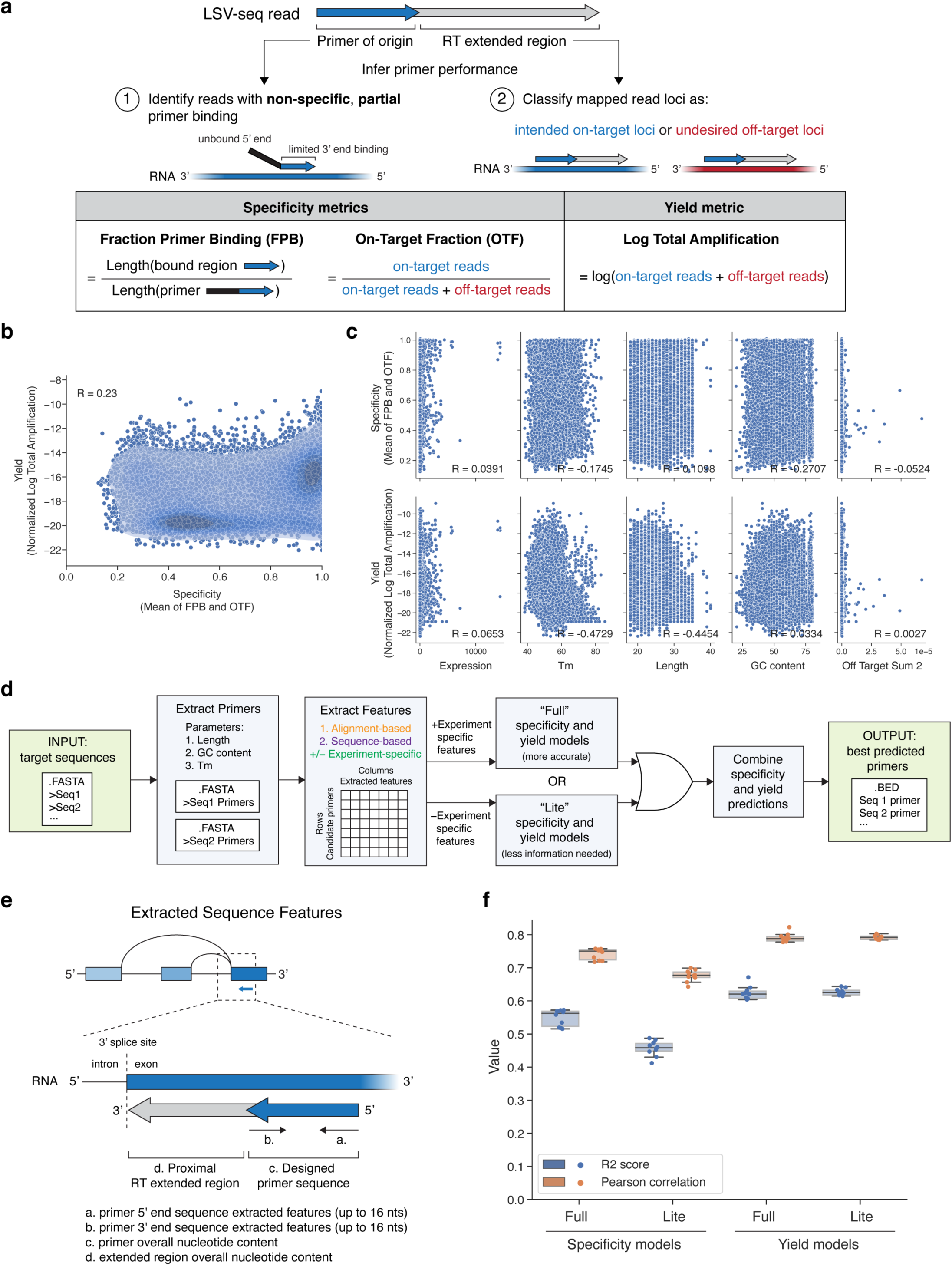
Primer selection pipeline based on Optimal Prime (OP) models. **a,** Overview of specificity and yield metrics derived to infer individual primer performance. **b,** Scatter plot depicting the correlation between the specificity metric (mean of FPB and OTF) and the yield metric (Normalized Log Total Amplification). Pearson correlation coefficient is given. **c,** Scatter plots depicting the correlation between various individual features thought to be important for determining primer performance and the specificity metric (mean of FPB and OTF) or the yield metric (Normalized Log Total Amplification). Pearson correlation coefficients are given and definitions of individual features are provided in Supplementary Table 2. **d,** Overview of processing steps in final primer selection pipeline. Pipeline allows use of either the “full” or “lite” specificity and yield models. **e,** Overview of encoded sequence-based features. Individual LSV-seq primer is shown as it would bind to the RNA during reverse transcription, along with the proximal region extended by the reverse transcriptase. **f,** Cross-validated performance of trained regression models. For each indicated model type, the calculated R2 score and Pearson correlation are given based on held-out 5-fold cross-validation splits conducted independently twice.

Since no simple combination of features was strongly predictive, we hypothesized that a more complex, dedicated machine learning model could significantly improve primer design (Fig. 2d). In order to build this model, which we named Optimal Prime (OP), we expanded the set of explanatory features from the initial 6 to over 1,000. We implemented our own primer design and feature extraction pipeline, dually inspired by the OligoMiner pipeline for RNA FISH probe design^17^ and prior feature extraction methods used for CRISPR-Cas9 guide prediction^18^. A ∼50-100 base pair target region is converted into all possible candidate primers by filtering for substrings satisfying relatively relaxed constraints for GC content, length, and melting temperature. The candidate primers are aligned with the BLAST alignment algorithm^19^ and on- and off-target alignments are passed into the NUPACK nucleic acid binding prediction algorithm^20^, as in previous work^17,21^, to create alignment-based features. An additional optional category of features includes those which are specific to the experimental context, such as gene expression levels for the targeted tissue or number of primers targeted to the same locus. Importantly, we also extracted hundreds of key sequence-based features for each candidate primer based on one-hot encodings of different positional nucleotide motifs (Fig. 2e). A full set of model features and their corresponding descriptions are given in Supplementary Table 2. The final set of features are passed into either “full” or “lite” specificity and yield models. The “full” model supplements this set of features with experiment-specific features to maximize prediction performance for LSV-seq. In contrast, the “lite” model is experiment-independent and therefore more flexible, potentially generalizing to other applications beyond LSV-seq. Finally, the separate predictions from the specificity and yield models are combined into a single score, allowing all candidate primers per locus to be ranked by predicted performance.

During model development, we tuned the design of our models through a combination of model selection, dataset filtering to remove noisy low-coverage data points and different formulations of the predicted variables (Supplementary Fig. S2). Interestingly, the boosted decision tree model exceeded the performance of the two deep learning-based architectures we benchmarked, including convolutional neural networks (CNNs) and transformer models (Supplementary Fig. S2a,b). Based on the mean performance across two independent 5-fold cross-validation procedures, we were able to generate highly accurate OP models based on the boosted decision trees for both specificity (full model Pearson’s R = 0.742 and R2 score = 0.549; lite model Pearson’s R = 0.676 and R2 score = 0.457) and yield (full model Pearson’s R = 0.822 and R2 score = 0.674; lite model Pearson’s R = 0.792 and R2 score = 0.627) (Fig. 2f). For the specificity model, the OP model represents a relative performance increase of 7.5-fold, out of a maximum possible of 13.7-fold, compared to the original maximum individual factor model performance of 7.3%.

### Validation and feature analysis of Optimal Prime models

While many prior primer design algorithms are either challenging to experimentally assess or are limited to relatively low-throughput validations, we leveraged the high-throughput scale of LSV-seq to directly validate the performance of our OP model at the bench across almost 1,000 new primers. We compared the performance of primers designed only considering classic heuristics and predicted on- and off-target alignments, versus primers newly re-designed based on our OP specificity model (Fig. 3a). Although these two sets of primers were designed to target the same 948 target LSVs, with one primer per target locus, we vastly improved the median specificity metric score from 0.55 in the non-OP design to 0.94 after incorporating our OP model (out of a maximum possible score of 1.0). We also greatly reduced the rate of primer dropout (primers with no detectable reads), from 8.9% (84/948 primers) to 0.3% (3/948 primers).

**Fig. 3.**
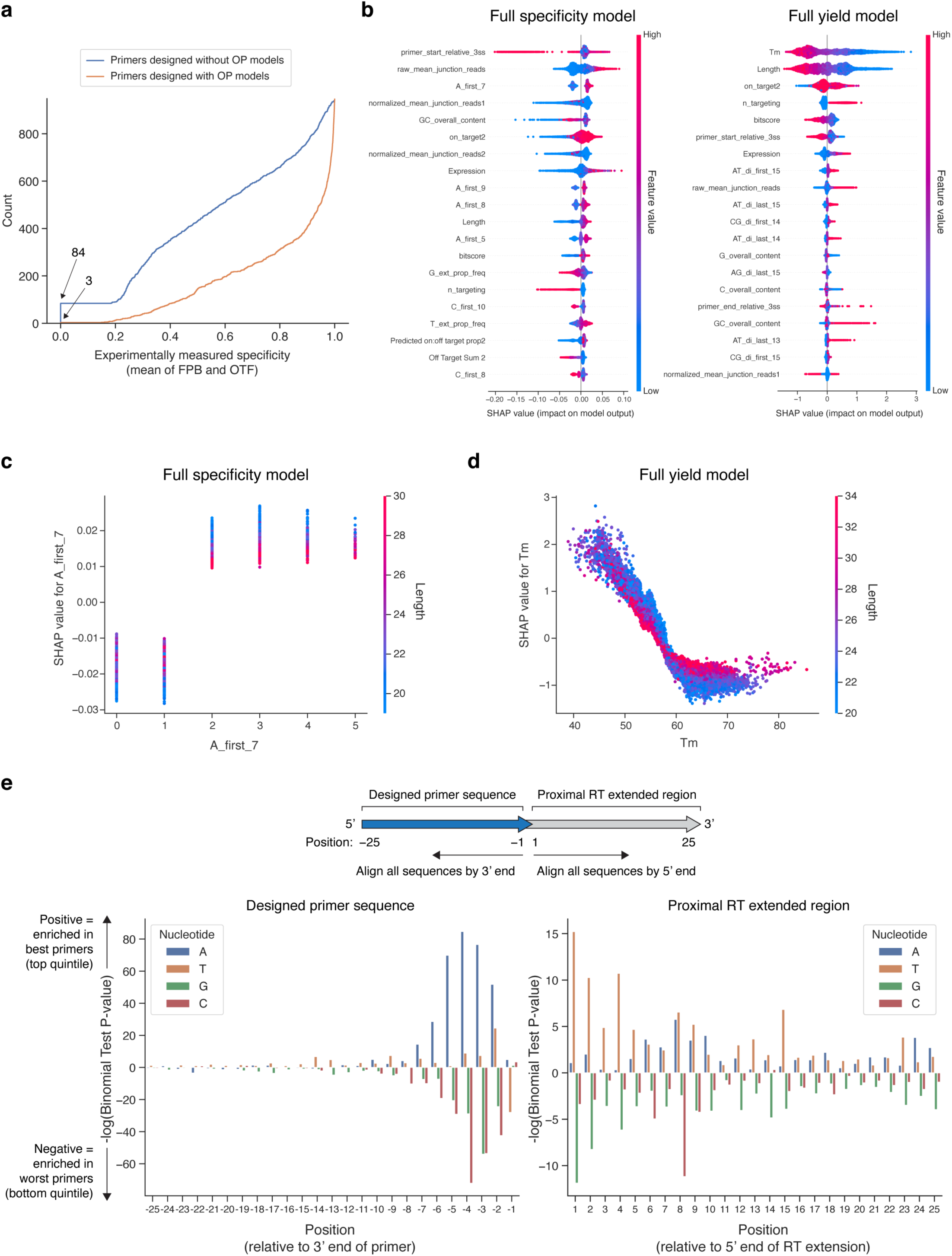
Experimental validation and interpretation of OP models. **a,** Cumulative distribution function for the performance of primers designed with (orange) or without (blue) the OP model, as measured by the specificity metric (mean of FPB and OTF), for the same set of target regions (n = 948). Number of primers with no observed value are noted by the arrowheads. **b,** Top features for full OP models. Features are ranked by mean absolute SHAP value magnitude. **c,d,** Informative interactions between top features for full models. For each feature indicated, its top most influential interactor by SHAP value magnitude is colored. **e,** Bar graph depicting the relative enrichment of specific nucleotides at the first 25 positions in both directions from the 3’ end of the primer, for either the best or lowest performance primer groups, as evaluated by the specificity metric (mean of FPB and OTF). For each position and nucleotide combination, a binomial test was conducted to determine the likelihood of the observed proportion in the top quintile of primers, compared to the proportion in the bottom quintile of primers.

Highlighting the explainability of the boosted decision tree model architecture, we interpreted the top features which contributed the most to primer performance via SHapley Additive exPlanation (SHAP) interaction values^22^. Although the same set of features are used in the specificity and yield models, their relative importance differs greatly (Fig. 3b). For the specificity model, we discovered a number of features which represent the adenosine content within the 7-9 nucleotides at the 3’ primer end. Conceptually, the 3’ end represents the nucleotides most important for formation of the reverse transcription initiation complex prior to elongation^23^. In contrast, for the yield model, the top model features were melting temperature and length, followed by a distinct set of sequence-based features. We further investigated more complex interactions between interpretable sets of top features per model. For the specificity model, a representative adenosine-rich 3’ end feature differentially modifies the impact on predicted performance for the length feature, in adenosine-poor primers (≤1 adenosine in 3’ end) compared to adenosine-rich primers (≥2 adenosines in 3’ end) (Fig. 3c). For the yield model, our analysis of the melting temperature and length features reveals that as primers grow longer, the relative effect of calculated melting temperature is somewhat decreased, represented in a shallower curve (Fig. 3d). Additionally, the point of inflection in determining model penalty for high melting temperature occurs close to the reverse transcription reaction temperature of 60°C, suggesting a link to real-world reaction conditions. Independently of our models, we also performed a binomial test for the relative enrichment of specific mononucleotide motifs in either the top-performing (top quintile) or bottom-performing (bottom quintile) primers for the specificity metric, as has been done in other nucleic acid prediction contexts^18^ (Fig. 3e). The most statistically significant differences are found in the increased preference for adenosines between nucleotides −1 and −7 from the 3’ end for top performing primers, which is consistent with their usage as top features in the OP specificity model. The guanine and thymine contents of the proximal extended region are also statistically significant determinants, and are likewise present within the top model features.

In order to increase the accessibility of our OP models to others for the design of LSV-seq primers, we implemented our primer selection pipeline as a webtool. We preliminarily investigated the effects of using the full versus lite models for the specific task of creating a relative ranking of primers, and noted an overall strong concordance in the rankings between the model types, although some specific points experience high discordance (Supplementary Fig. S3b,c). Thus, we decided to allow for the selection of two distinct run modes, both of which bypass the most computationally expensive steps and greatly reduce the runtime on a live web platform. “Lite Mode”, which implements our lite OP models, allows a more flexible array of inputs, based on either a BED6 file specifying mouse or human chromosome coordinates, a FASTA file, or a selection from a list of LSVs. “Full Mode” implements our more accurate full OP models. To achieve this, we ran MAJIQ across 54 GTEx tissues, allowing us to comprehensively catalog all >190,000 target LSVs we could detect across all human tissues. We then exhaustively generated >16 million total candidate primers from these identified target LSVs, and precomputed the BLAST alignment and NUPACK binding prediction steps, which took ∼7 days on our cluster. To run the webtool in “Full Mode”, the user selects from a list of LSVs and either specifies preloaded expression values for a given tissue of interest, or supplies their own list. For both run modes, the final output is a list of the top 10 primer sequences for each region and their corresponding prediction scores.

### Benchmarking and validation of LSV-seq

To benchmark LSV-seq, we used Jurkat T-cells which we have previously shown undergo reproducible splicing changes upon stimulation with phorbol myristyl acetate (PMA)^24^. Having established high capture rates for our optimized primers, we collected stimulated and unstimulated Jurkat cells in matched biological triplicate and then used the same RNA to generate either standard RNA-seq libraries (sequenced to a minimum depth of 100 million reads) or LSV-seq libraries with a pool of 1,991 targeting primers designed using our Optimal Prime pipeline (sequenced to a depth of 10 million reads). LSV-seq libraries were highly specific, with more than 95% of reads corresponding to targeted regions, while the same regions were covered by only ∼1-2% of reads in the RNA-seq dataset (Fig. 4a). We also achieved a median of ∼19 and mean of ∼230 fold-enrichment overall (Fig. 4b).

**Fig. 4.**
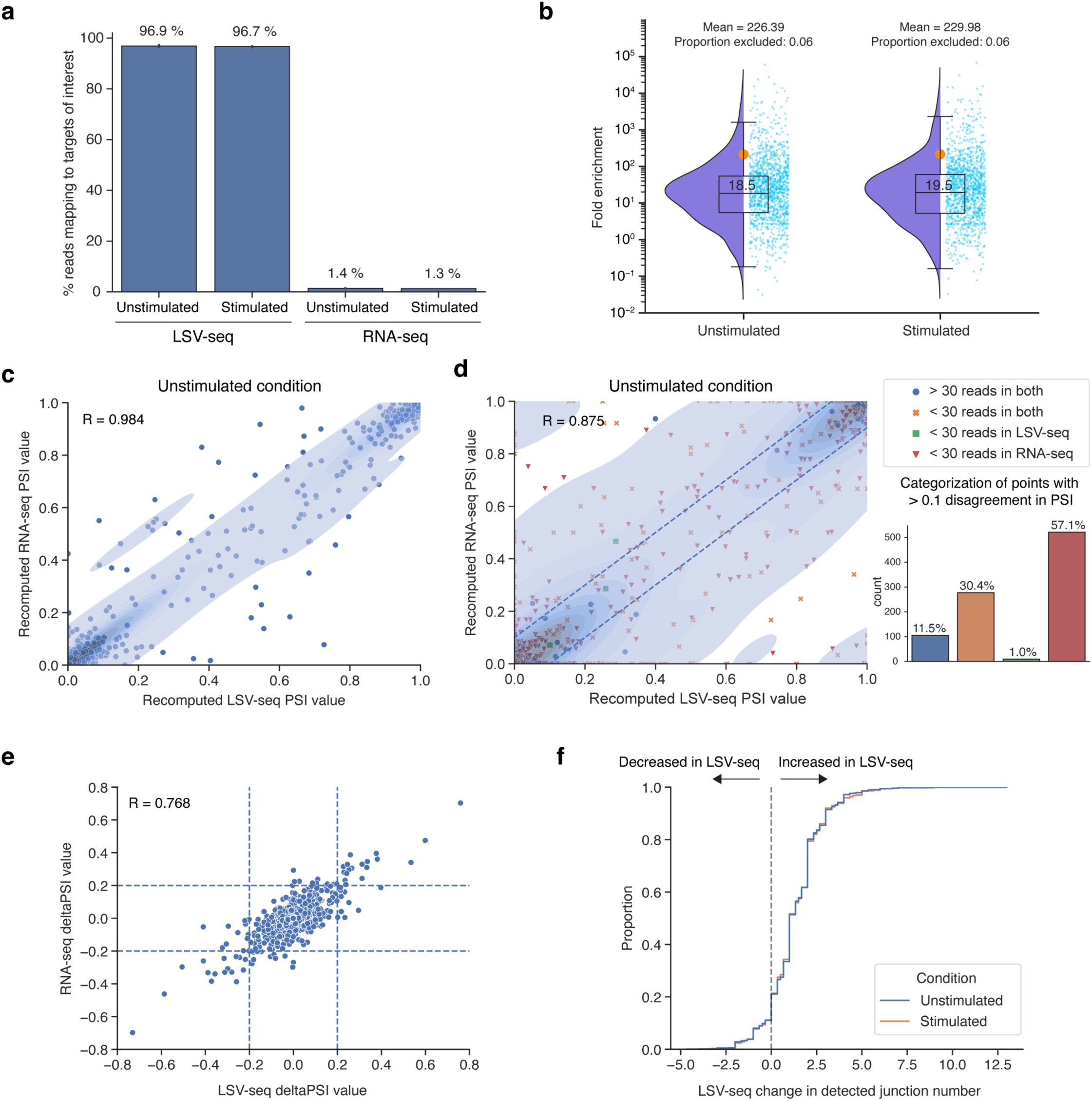
LSV-seq recapitulates gold standard RNA-seq measurements and enriches low-coverage RNA-seq measurements. **a,** Percent of LSV-seq reads mapping to targeted splicing events of interest (n = 1991) in Jurkat T-cells. Also shown are the percent of reads mapping to the same splicing events in RNA-seq data. **b,** Raindrop plot depicting the mean LSV-seq fold enrichment over RNA-seq for unstimulated (n=3) and stimulated (n=3) Jurkat T-cell biological replicates per splicing event. The box plot markings represent the 0th to 100th percentiles in increments of 25, with the median marked at the 50th percentile, and the mean denoted as an orange dot. A small proportion of targeted events which did not reach at least 5 detected reads in either LSV-seq or RNA-seq were excluded from analysis, as indicated above the plots. **c,** Scatter plot comparing the PSI values for the same splice junctions in either saturated LSV-seq or RNA-seq datasets (for the unstimulated Jurkat T-cell condition), across only splicing events with at least 30 mean reads of coverage in both. Each splicing event is reduced to one splice junction selected at random. Pearson correlation coefficient is given. **d,** Scatter plot comparing the PSI values for the same splice junctions in equally downsampled LSV-seq or RNA-seq datasets (for the unstimulated Jurkat T-cell condition), across all splicing events. Only junctions observed in both LSV-seq and RNA-seq are considered when calculating the PSI value. Each splicing event is reduced to one splice junction selected at random. Points which have high disagreement are defined as those having over 0.1 PSI discordance (consisting of points lying outside the dashed lines). The bar plot quantifies the coverage categories of these discordant points. Pearson correlation coefficient is given. **e,** Scatter plot comparing the stimulated - unstimulated deltaPSI values for the same splice junctions in full-depth LSV-seq or RNA-seq datasets, across splicing events with at least 50 reads of coverage in both. All junctions, whether or not they are observed in both, are considered in calculating the deltaPSI values. Dashed lines indicate significant deltaPSI values over 0.2. Pearson’s correlation was calculated by shrinking all points within 0.2 deltaPSI (within the inner box) to 0. **f,** Cumulative distribution functions plotting the mean difference in number of junctions detected in equally downsampled LSV-seq and RNA-seq. Positive values indicate more junctions detected in LSV-seq, while negative values indicate more junctions detected in RNA-seq.

One key characteristic of the LSV-seq method presented here is its tight integration with MAJIQ. As discussed earlier, this integration is reflected in the type of AS events LSV-seq inherently captures at the experimental level, which correspond to target LSV by MAJIQ’s formulation. Practically, during post-sequencing data processing, this also requires quantification using the MAJIQ PSI/deltaPSI algorithms and visualization using the VOILA package, which were all originally designed for standard RNA-seq data. To accomplish this, we created a new independent python package and several compatibility updates within the base MAJIQ algorithm to allow us to seamlessly analyze splicing events for LSV-seq. To fairly compare the resulting RNA-seq and LSV-seq quantifications, we performed analyses of both full-depth datasets and data downsampled to equal read depths (Supplementary Fig. S4a). First examining only splicing events with high coverage in both full-depth LSV-seq and RNA-seq (defined as having at least 30 reads per event), we noted near identical PSI values (R = 0.984), demonstrating that LSV-seq provides quantifications that are highly consistent with those from high-coverage RNA-seq (Fig. 4c). To further assess the reproducibility between LSV-seq and RNA-seq, we produced Bland-Altman plots, which are commonly used to compare the agreement between two different assays, by plotting the mean of the assays (mean of LSV-seq and RNA-seq PSI values) against the magnitude of disagreement (difference between LSV-seq and RNA-seq PSI values) (Supplementary Fig. S5a)^25^. The limits of agreement between LSV-seq and RNA-seq are represented by the interval from −0.15 to 0.14, suggesting strong concordance. Comparatively, the limits of agreement are −0.08 to 0.08 for LSV-seq and RNA-seq samples internally (Supplementary Fig. S5b,c).

Next, when we removed our read threshold filter and instead looked at splicing events across equally downsampled LSV-seq and RNA-seq, the correlation dropped considerably (R = 0.875) (Fig. 4d). PSI values which differed by more than 0.1 between LSV-seq and RNA-seq could represent technical issues with either LSV-seq, RNA-seq, or both. Our analysis of these disagreeing quantifications categorized a large proportion of such splicing events as having low coverage (<30 reads) in only RNA-seq, while effectively none of these events have low coverage (<30 reads) in only LSV-seq. This suggests that a majority of quantifications that disagree between the two assays are likely due to coverage issues with RNA-seq specifically. In addition to calculating PSI values across replicates, we also examined our ability to accurately profile the difference in mean PSI values, or deltaPSI, between the stimulated and unstimulated condition groups, using the same 30 read minimum threshold for analyzed events as in the previous analysis (Fig. 4e). Notably, since this analysis requires quantifying events in two conditions, we included junctions reported as low coverage in either assay technology. Even with this addition, we still observed a strong correlation between deltaPSI values from RNA-seq and LSV-seq (R = 0.768). Reassuringly, the deltaPSI values measured in LSV-seq between matched pairs have significantly lower variance across read coverage bins compared to RNA-seq (Supplementary Fig. S4b,c).

As another key benchmark, we assessed the ability of LSV-seq to consistently capture splice junctions in each quantified splicing event. For equally downsampled LSV-seq and RNA-seq datasets, we compared the difference in the number of splice junctions detected. For ∼70% of splicing events within a biological condition, LSV-seq detects, on average, at least one extra splice junction per LSV compared to RNA-seq (Fig. 4f). Moreover, the vast majority of these splice junctions are likely true biological occurrences, as 8406/8521 (98.7%) of them can be detected in the full-depth RNA-seq dataset (Supplementary Fig. S4d).

Finally, we assessed the ability of LSV-seq to approximate changes in expression. Although LSV-seq was primarily optimized to quantify PSI and deltaPSI values, the log2 fold-changes in per-primer LSV-seq coverage can potentially be helpful as an orthogonal metric to discriminate between tissue conditions. We compared the mean LSV-seq log2 fold-changes in single-primer coverage to the mean of RNA-seq log2 fold-changes in whole-gene summed transcripts per million (TPM) values. While we do not necessarily expect LSV-seq coverage of a single targeted ∼150-base region to directly correspond to RNA-seq whole-gene expression, we observed an unexpectedly strong correlation between LSV-seq and RNA-seq in measured coverage differences (R = 0.922) (Supplementary Fig. S4e), suggesting LSV-seq could be used to simultaneously track changes in target coverage. Moreover, the strength of the LSV-seq/RNA-seq correlation is similar to the strength of correlation across true biological replicate measurements (ranging from R=0.967 to R=0.971).

### LSV-seq recovers previously uncharacterized tissue-specific splicing events in GTEx

Next, we reasoned that the enhanced sensitivity of LSV-seq could recover previously unquantifiable, low-coverage splicing events. We first set out to assess how many of the AS events detected in existing large datasets such as GTEx may suffer from limited quantifiability. In representative tissue-wide GTEx RNA-seq datasets we analyzed using our MAJIQ algorithm^26^, we found that only an average ∼10% of reads span splice junctions, preventing consistent capture of less abundant isoforms^27^ (Fig. 5a). Because of the sparse coverage of splice junctions, up to ∼44% of AS events we detected in the GTEx data had 25 or fewer mean reads of coverage and could not be reliably quantified in a significant fraction of samples (where “reliable” is defined by the LSV having at least 10 reads) (Fig. 5b and Supplementary Fig. S6a). Moreover, the large majority of these difficult-to-quantify AS events reside in well-expressed genes (with at least 5 TPM) (Fig. 5c), hinting at their potential biological relevance.

**Fig. 5.**
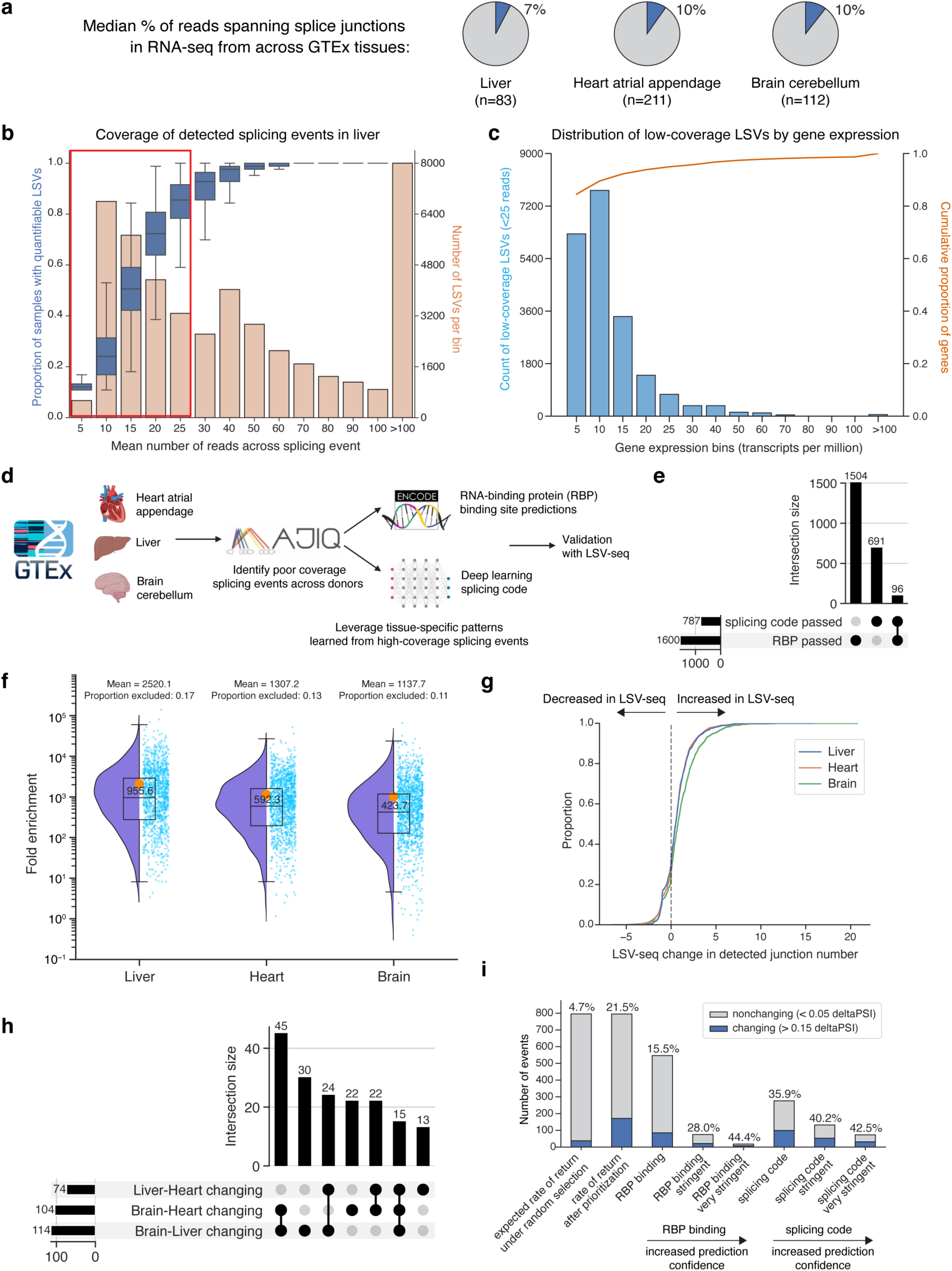
LSV-seq captures splicing events that are unquantifiable in GTEx RNA-seq datasets. **a,** Pie charts illustrating low capture rate of splice junction reads in GTEx RNA-seq datasets across 3 tissues. **b,** For the GTEx liver dataset, histogram in orange illustrating distribution of read coverage for all detectable LSVs, overlaid with box plot in blue illustrating decrease in quantifiability rate at low read coverage. Red box highlights the ∼44% of AS events with 25 or fewer mean reads of coverage that cannot be reliably quantified. **c,** For the GTEx liver dataset, histogram in light blue illustrating the distribution of gene expression by TPM for low-coverage LSVs, overlaid with the cumulative distribution function in orange for all expressed genes. **d,** Overview of the pipeline used to prioritize low-coverage splicing events which are predicted to be tissue-specific across 3 different GTEx tissues. **e,** Upset plot depicting the relative overlap of the initial list of candidate events identified by each pipeline. **f,** Raindrop plot of the mean fold-enrichment of LSV-seq over RNA-seq for the mean splicing event coverage over 3 technical replicates, compared to the mean coverage over the entire GTEx dataset for each tissue. For each technology, read coverage is normalized by the mean library size. **g,** Cumulative distribution functions plotting the difference in the number of mean junctions per splicing event detected across full-depth LSV-seq replicates, compared to the full-depth GTEx RNA-seq dataset, per tissue. Positive values indicate more junctions detected in LSV-seq, while negative values indicate more junctions detected in RNA-seq. **h,** Upset plot categorizing changing events captured by LSV-seq. For each changing LSV, we noted which specific pairwise tissue comparison(s) they were changing between. **i,** Summary of LSV-seq validation results for the splicing event prioritization pipeline. Different subcategories of events, based on their original prioritization pipeline and prediction confidence, are shown with their relative enrichment for confidently changing events (> 0.15 deltaPSI in at least 1 pairwise comparison), out of total confidently changing and nonchanging (< 0.05 deltaPSI in all pairwise comparisons) events.

After establishing that the GTEx RNA-seq data did indeed have a large number of splicing ‘blind spots’, we devised a strategy to prioritize which of these events to target with LSV-seq (Fig. 5d and Supplementary Fig. S6). Specifically, we aimed to recover tissue-specific splicing events which are changing in at least 1 of 3 pairwise tissue comparisons between liver, heart atrial appendage and brain cerebellum. Based on our initial estimates, targeting splicing events at random would identify tissue-specific events at an unacceptably low rate of 4.7%. To mitigate this, we created a prioritization pipeline to nominate putative tissue-specific splicing events that we could then experimentally validate with LSV-seq. We first identified all unquantifiable low-coverage LSVs (<25 mean reads of coverage) across the 3 selected tissues. We then classified each event as putatively differentially spliced between tissues based on two separate prediction pipelines. The first uses the Encyclopedia of DNA Elements (ENCODE) CLIP-seq data that captures the binding sites of RNA binding proteins (RBPs)^28,29^, while the second uses an in-house splicing code deep learning model. Surprisingly, these two pipelines predicted largely non-overlapping sets of putative tissue-specific splicing events (Fig. 5e).

We then used our OP primer design pipeline to create a pool of 1,514 primers capturing 1,400 unique targets and performed LSV-seq on RNA from human liver, heart atrial appendage, and brain cerebellum. LSV-seq was able to greatly boost the coverage of targeted splicing events between a median of 424- to 956-fold depending on the tissue, far exceeding the original GTEx RNA-seq coverage level (Fig. 5f). Due to the corresponding increase in consistency of junction-level coverage, we also noted a large gain in the number of detected junctions per event (Fig. 5g). When we assessed the deltaPSI values we captured between each pairwise tissue comparison, we found we were able to call 292 unique pairwise differences (deltaPSI>0.15), corresponding to 171 unique LSVs (Fig. 5h), with the most differences found between brain cerebellum and either of the other two tissues. Our overall rate of return was 21.5% for tissue-specific events, over 5 times the rate that would be expected with random selection (Fig. 5i). Our splicing code pipeline in particular performed especially well, with 35.9% of candidate events being validated as truly changing between tissues. In contrast, the RBP-based pipeline nominated twice as many candidate events yet returned a similar number of truly changing events (15.5% true positive rate), demonstrating that the false positive rate of the splicing code pipeline is much lower.

To validate our original tissue-specific prediction pipeline, we assessed if the *post hoc* tissue-specific splicing change frequencies we empirically observed in LSV-seq reflected our *a priori* knowledge of different subcategories we expected to be informative. If our RBP binding and splicing code pipelines performed correctly, then we expect that as we increase the stringency of the threshold for calling putative changing events, the rate of false positives should decrease, at the cost of total true positives detected. Indeed, for both pipelines, we observe exactly this, with fewer false positives called as we increased the prediction threshold stringency in each pipeline, reaching a maximum validation rate of 44.4% in the RBP binding pipeline and 42.5% in the splicing code pipeline. For each tissue-specific splicing regulatory RBP we identified in our RBP binding pipeline, we also calculated the expected discovery rate based on the prevalence of the RBP’s binding sites within tissue-specific splicing events in high-coverage RNA-seq. We then explicitly tested the concordance between each RBP’s expected discovery rate from RNA-seq and its true observed discovery rate in LSV-seq, and observed a strong positive correlation (R = 0.75) (Supplementary Fig. S6e).

Next, we further examined the 171 tissue-specific splicing events we had newly recovered using LSV-seq and which previously could not be captured by the limited coverage of the GTEx RNA-seq data. Within this set, we found many examples of splicing that were highly specific to each of the three tissues assayed (Supplementary Fig. S7), often involving splice junctions that were detected in only one tissue type. Among the splicing events we found to be unique to the brain cerebellum, we achieved robust splicing detection of an *ENAH* microexon (Supplementary Fig. S7a), that was previously identified in a study of neuronal microexons^30^. Our LSV-seq results for the *ENAH* microexon (PSI = 0.39) were consistent with this independent microexon study (cerebellum PSI = 0.34), supporting the accuracy of LSV-seq in quantification of targeted events. Another brain-specific alternative splicing event involved the *RAB3GAP1* gene. Rare genetic variants within the *RAB3GAP1* gene have previously been reported to cause the Micro and Martsolf autosomal recessive disorders, both of which are associated with a collection of profound neurological deficits, including visual impairment, brain abnormalities, and hypotonia^31^. One such pathogenic mutation induces a frameshift very early within the N-terminal domain, likely almost fully abrogating the transcript, leading to the suggestion that a previously detected alternatively spliced isoform lacking the first 50 N-terminal amino acids could partially rescue RAB3GAP1 function. Using LSV-seq, we measured the relative presence of this alternatively spliced isoform (blue junction) compared to the dominant isoform (red junction) across tissues, whereas previous GTEx RNA-seq coverage was insufficient for detection (Fig. 6a). We found that the alternatively spliced isoform is expressed specifically in the neural tissue we tested, brain cerebellum, compared to the other two tissues, heart atrial appendage and liver (Fig. 6b). Interestingly, this change in alternative splicing corresponds with the presence of binding sites for RBPs TIA1 and KHSRP (Fig. 6a), which both have significantly higher expression in the brain cerebellum compared to the other tissues (Fig. 6c).

**Fig. 6.**
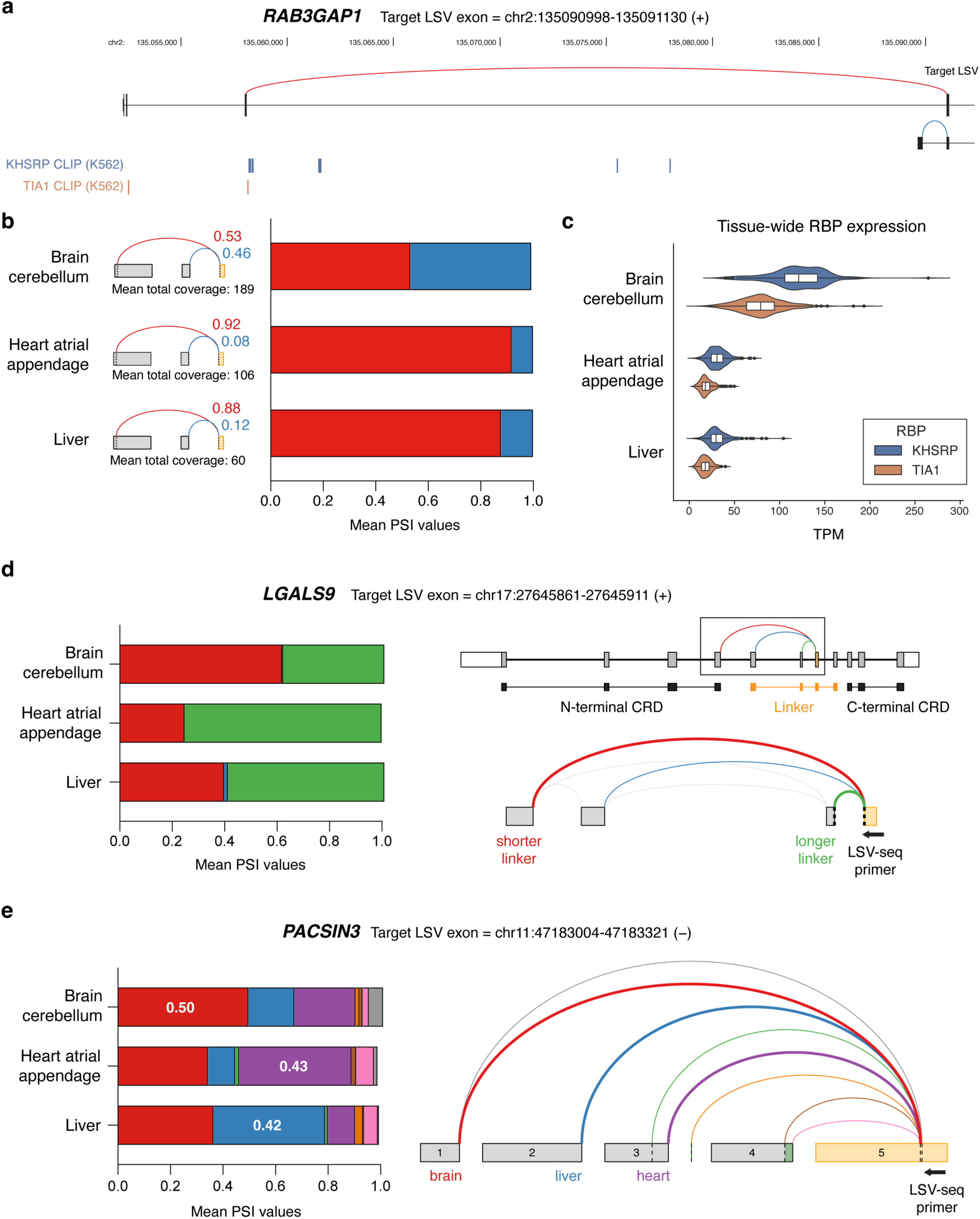
Tissue-specific splicing event in *RAB3GAP1* recovered by LSV-seq. **a**, *RAB3GAP1* gene track with splice junctions shown as red or blue arcs. Also shown below are ENCODE CLIP IDR binding peaks for the RBPs KHSRP and TIA1. **b**, *RAB3GAP1* LSV splicing and PSI values across tissues as measured by LSV-seq. **c**, Expression of KHSRP and TIA1 RBPs across all samples in GTEx for each of the three tissues. **d,** *LGALS9* LSV splicing and PSI values across tissues as measured by LSV-seq. Also shown are the gene structure of *LGALS9* with annotation of the N/C-terminal CRDs and alternatively spliced linker region, and the Voila splice graph of the LSV (corresponding to boxed area). **e,** Complex LSV splicing and PSI values for *PACSIN3* across tissues as measured by LSV-seq. Also shown is the Voila splice graph with the dominant splice junctions for each tissue represented by thicker arcs. Numbers in PSI graph indicate PSI value for that specific splice junction.

In addition to events that were unique to a single tissue, we also recovered tissue-specific splicing that was quantitatively different across all three tissues, such as in *LGALS9* (Fig. 6d). *LGALS9*, or galectin-9, plays an important role in immunomodulation through binding to the TIM-3 receptor on immune cells^32^. As this interaction can suppress immune responses, the galectin-9/TIM-3 axis is being actively investigated as a potential target for immunotherapy. Galectin-9 is expressed as several different isoforms that differ in the length of the linker region connecting its two carbohydrate recognition domains (CRDs). Differences in linker length affect the rotational freedom of the CRDs and multivalency of galectin-9, which can then impact its interactions with other proteins^33^. Using LSV-seq, we detected differential splicing in this linker region, with heart tissue favoring galectin-9 isoforms containing a longer linker and brain tissue favoring isoforms with a shorter linker (Fig. 6d). Finally, LSV-seq also captured complex splicing in many cases, highlighting its unique ability to detect multiple known and de novo junctions at each targeted LSV using only a single primer. For example, we detected particularly complex splicing in the 5’ region of *PACSIN3*, with many different splice junctions being used across all three tissues. Interestingly, LSV-seq also revealed that each tissue exhibited dominant splicing of a different junction, suggesting tissue-specific regulation of the *PACSIN3* 5’UTR (Fig. 6e). Altogether, our results illustrate the ability of LSV-seq to capture quantifications for biologically relevant splicing events that may escape detection with conventional RNA-seq.

## DISCUSSION

Although short-read RNA-seq continues to be the standard approach for splicing analysis, it is inefficient at capturing the splice junctions needed for accurate quantification. Indeed, our analysis of the GTEx RNA-seq dataset revealed inadequate detection of a sizable fraction of biologically important splicing variation across tissues. To address this limitation, we developed LSV-seq, a method for targeted detection and quantification of up to thousands of splicing events of interest. In comparison to conventional RNA-seq, we demonstrate that LSV-seq provides significantly improved coverage at targeted events and accurate splicing quantification even with markedly reduced sequencing depths. We also find that LSV-seq can recover splicing information at events with poor coverage in GTEx RNA-seq data to reveal novel forms of tissue-specific splicing. Altogether, LSV-seq offers an efficient and versatile method for the study of AS in humans and other organisms with comparably complex transcriptomes.

LSV-seq is inspired by a previous method, MPE-seq, that was originally developed for detection of splicing in yeast^11^. Like LSV-seq, MPE-seq enriches for RNA regions of interest by performing multiplexed RT with pools of target-specific primers. However, our initial attempts to apply MPE-seq to human RNA resulted in a majority of off-target reads (Supplementary Fig. S1). Additionally, we note that a previous study performed primer extension based on MPE-seq in human cells, although only in a low-throughput format for a single intron^34^. We reasoned that our observations could be due to the much greater complexity of the human transcriptome (∼250,000 annotated transcripts^35^) as compared to yeast (∼7,000 annotated transcripts^36^). As a result, we focused most of our efforts on improving the specificity of the RT reaction through both experimental and computational optimizations. For example, we incorporated a linear amplification step, used in some single-cell RNA-seq protocols, allowing us to reduce the amount of input RNA needed by at least ten-fold. However, as the 5 ug of input RNA we recommend may still exceed what is available, especially in the context of valuable clinical samples, further experiments will be needed to determine the compatibility of LSV-seq with lower input conditions.

When compared to other targeted sequencing methods such as CaptureSeq^6^, RASL-Seq^8^, and TempO-seq^9^, one key advantage of LSV-seq is its ability to detect all junctions adjacent to the targeted region using only a single primer. Thus, with far fewer primers, LSV-seq can capture both simple and complex splicing events, including those with novel, unannotated junctions. Although each LSV-seq primer is limited to detecting splicing variation at target LSVs, we found that this is sufficient to capture ∼80% of splice junctions (Fig. 1c). We anticipate that variation at the remaining junctions found in source LSVs could be captured by using combinations of primers anchored near each downstream junction, although this remains to be tested. More recently, long-read technologies have emerged with the ability to sequence full-length transcripts. Despite this advantage, current long-read RNA-seq studies are generally better equipped to tackle novel isoform detection, while accurate quantification remains an active area of development. Future work marrying LSV-seq primer pools with long-read sequencing could facilitate sensitive and accurate splicing quantifications, analogous to what we have shown for short-read RNA-seq.

We credit the Optimal Prime machine learning models for the strong performance of LSV-seq. Early in the development of our method, we discovered a lack of tools suitable for the design of RT-specific primers and noticed that most related protocols repurposed pipelines originally created for other multiplexed assays such as microarray or fluorescence in situ hybridization. Indeed, our first attempts at multiplexed RT using these off-the-shelf tools resulted in primers with significant off-target behavior or failure to prime desired targets. We also reviewed other tools from the literature tailored for designing splice junction-spanning primers and found these were also not well-suited for LSV-seq, either because they lacked adequate experimental validation or focused on generating pairs of PCR primers rather than target-specific RT primers^37–40^. Thus, we sought to use a data-driven approach to first discover optimal RT primer properties and then train machine learning models to predict primer performance. For this, we defined two optimal priming measures, namely target specificity and yield, and found that none of the classical primer features, such as Tm or GC content, correlated well enough on their own with these measures to be used independently. However, by combining features together using the OP algorithm, we were able to substantially improve both the yield and specificity of the designed primers. Interestingly, models based on boosted decision trees outperformed both convolutional neural networks and transformers, suggesting that our handcrafted features provide near optimal design for our current dataset. However, we anticipate the deep learning models could eventually overtake the performance of our boosted decision tree models when given more training data from additional LSV-seq experiments.

One important observation regarding the primer optimization is that the OP lite model, which ignored off-target transcriptomic alignments and relative gene expression, still offered good performance compared to the OP full model. This lite model thus offers greater flexibility when condition-specific measurements are not available and for applications beyond LSV-seq. To enable the research community to take advantage of our OP algorithm for primer design, we have made it accessible via a webtool. By precomputing computationally expensive values or by running an alignment-free version, the webtool offers lightning-fast retrieval of optimal primers for targets of interest.

Our work also demonstrates the strengths and limitations of current approaches for the prediction of human tissue-specific splicing events. Here, one approach used was the incorporation of known RBP data^41,42^ while the other was based on sequence context and tissue identity^42–44^. The first approach was represented by a pipeline for analysis of ENCODE CLIP data, while the second was based on a tissue-specific splicing code model (without CLIP information). Our results using these two approaches reveal that there is little concordance in the predictions provided by our RBP binding and splicing code pipelines. One possible explanation for this result is that the splicing code pipeline focuses on binary cassette event prediction, while the RBP binding pipeline allows for prediction across a wider range of splicing event types. The lack of overlap may also indicate that our current splicing code model does not yet fully reflect the underlying biology of RBPs and their corresponding binding site sequence motifs. Although there is still room for improvement, the splicing code pipeline had a far lower false-positive rate for discovery of tissue-specific splicing events compared to the RBP binding pipeline. Thus, our analysis validates the concurrent use of both pipelines in order to maximize the number of tissue-specific changing events we can recover. Future work can also further explore combined usage of splicing code modeling with data from RBP binding assays, as was done previously^41,42^. Regardless of the exact model or pipeline, LSV-seq provides an efficient method to validate and further refine such computational splicing predictions.

From our prioritized selection of targets that had limited coverage in GTEx RNA-seq data, we recovered 171 events with tissue-specific splicing using LSV-seq. Across the three tissues we assayed (brain cerebellum, heart atrial appendage and liver), we discovered several instances of splicing that were highly specific to each tissue type, as well as splicing that was less specific but quantifiably different between tissues. While we highlighted examples in *RAB3GAP1, LGALS9* and *PACSIN3*, the majority of the splicing variation we detected is uncharacterized and our results nominate splicing events of interest for further functional analysis. Thus, LSV-seq provides the ability to focus in unexplored areas to recover splicing information and generate novel hypotheses about isoform regulation and function. Our findings also motivate further work to improve the mapping of alternative splicing across human tissues.

We anticipate that LSV-seq could be directly transferred in its current form to a number of additional applications. For instance, it has the potential to recover extremely rare splice junctions which are especially challenging to detect even with very deep RNA-seq. Targeted sequencing was recently used to recover transient splicing intermediates at putative recursive splicing sites in human cells, but was only performed for a single intron^34^. In addition to its sensitivity, LSV-seq is much more cost-effective than RNA-seq and could make it more feasible to study splicing across large numbers of samples or conditions. For instance, LSV-seq could be used in conjunction with high-throughput drug or genetic perturbation to quantify effects on thousands of AS events across thousands of different conditions. Such high-dimensional ‘many-by-many’ experiments could provide valuable insight into the complex network underlying regulation of AS. While larger LSV-seq primer pools do add an appreciable upfront reagent cost, each pool can be used for thousands of samples, and as experiments scale up in size, this primer cost is significantly offset by the 5-10 fold reduction in sequencing depth that is needed for LSV-seq. Although our study used at most ∼2000 primers, we expect LSV-seq to scale well to larger numbers of targets. However, the actual limits of LSV-seq remain unknown and we predict that increasing the number of targeted events will likely lead to more off-target priming as well. Our primer selection pipeline could also be broadly useful for various other contexts which require region-specific RT primers. For instance, due to the stringent requirement for >5000 reads per nucleotide within analyzed transcripts, SHAPE-Seq relies on numerous RT primers tiled across specific genes of interest^45,46^. For SHAPE-Seq and other related methods that analyze RNA structure, our primer selection pipeline could directly improve assay throughput and performance.

In summary, we have developed LSV-seq as a sensitive and cost-effective method for the detection and quantification of alternative splicing. We envision that LSV-seq will also enable studies in additional areas of RNA biology, helping to better capture other challenging yet important features of the transcriptome.

## METHODS

### Cell culture

The clonal Jurkat T-cell line (JSL1) was cultured and stimulated as previously described^47^. Cells were maintained in RPMI 1640 media supplemented with 5% fetal bovine serum, 100 U/mL penicillin and 100 ug/mL streptomycin. For stimulation, cells were seeded at 4 x 10^5^ cells/mL and treated with 20 ng/mL of the phorbol ester PMA (524400, Millipore Sigma). A separate culture of unstimulated cells was seeded at 2.5 x 10^5^ cells/mL. After 48 hours, successful stimulation was confirmed by staining cells for the activation marker CD69 (310905, Biolegend) followed by flow cytometric analysis (data not shown). Cells were then collected and flash frozen on liquid nitrogen in aliquots of 5×10^6^ cells. To generate 3 total biological replicates, PMA stimulation was done on independent days.

### RNA isolation and processing

Total RNA was purified from cell pellets with the Maxwell RSC 48 instrument using the Maxwell RSC simplyRNA Cells Kit (AS1340, Promega). Poly-A selection was subsequently performed using the NEBNext Poly(A) mRNA Magnetic Isolation Module (E7490L, New England Biolabs). LSV-seq was performed as described below. RNA-seq library preparation and paired-end 150 bp sequencing was performed by Novogene. Briefly, mRNA was purified from total RNA using poly-T oligo-attached magnetic beads. After fragmentation, first strand cDNA was synthesized using random hexamer primers. Then the second strand cDNA was synthesized using dUTP. The directional library was ready after end repair, A-tailing, adapter ligation, size selection, USER enzyme digestion, PCR amplification, and purification. After quality control, libraries were pooled and sequenced on the Illumina platform to a minimum depth of 100 million reads per library. Total RNA from human adult normal tissues (heart right atrium, liver, and brain cerebellum) was obtained from a commercial vendor (BioChain Institute Inc.). For each tissue, 3 technical replicates of LSV-seq were performed as described below.

### Preparation of MPE-seq libraries

MPE-seq libraries were prepared as previously described with minimal modification^11,13^. The 50- and 381-primer pools used for MPE-seq were designed while attempting to minimize predicted off-target alignments and basic considerations such as GC content and melting temperature, without the use of the Optimal Prime primer selection pipeline. On and off-target read percentages were calculated as described below for analysis of LSV-seq data.

### Preparation of LSV-seq libraries

#### First-strand synthesis reaction

Primer pools were designed according to our primer selection pipeline and ordered as a single oligo pool of hundreds to thousands of primers (oPools, Integrated DNA Technologies), see Supplementary Table 5 for primer sequences. 5 ug of total RNA was used directly in the reaction or following optional poly-A purification with the NEBNext Poly(A) mRNA Magnetic Isolation Module (NEB, E7490S). RNA was mixed with 2.5 uL of the primer pool (diluted to 400 pM/oligo) in a total volume of 20 uL, also containing a final concentration of 2 mM of each dNTP (N0447S, NEB) and 1X Maxima buffer (EP0752, Thermo Scientific). To denature the RNA and allow the primers to hybridize to the target RNA with high specificity, the reaction was incubated on a thermal cycler in a touchdown cycle from 85°C to 60°C, decreasing 1°C per minute for 25 minutes total, followed by a hold at 60°C. While maintaining the primer-RNA mixture at 60°C, 20 uL of reverse transcription mixture was directly added, consisting of 1 uL of 200 U/uL Maxima H Minus Reverse Transcriptase (EP0752, Thermo Scientific), 1 uL of 5 U/uL ThermaStop-RT reagent (TSTOPRT, Sigma Aldrich), 1 uL of 100 mM RNaseOUT Recombinant Ribonuclease Inhibitor (10777019, Invitrogen) and 1X Maxima buffer. The reverse transcription reaction was incubated at 60°C for 80 minutes, followed by heat inactivation at 85°C for 5 minutes. The reaction was then cleaned up with 40 uL of RNAClean XP beads (A63987, Beckman Coulter) and resuspended in 40 uL of water.

#### Second-strand synthesis reaction

To the 40 uL purified first-strand reaction, we added 40 uL of second-strand synthesis reaction mixture containing a final concentration of 0.2 mM of each dNTP, 0.55 uL of 10 U/uL E. coli DNA Ligase (M0205L, NEB), 2.08 uL of 10 U/uL E. coli DNA Polymerase I (18010025, Invitrogen), 0.55 uL of 5 U/uL E. coli RNase H (M0297L, NEB), and 1X NEBNext Second Strand Synthesis Reaction Buffer (B6117S, NEB). The reaction was incubated for 2 hours at 16°C, cleaned up with 80 uL of RNAClean XP beads, and resuspended in 10 uL of water.

#### In vitro transcription reaction

To the 10 uL purified second-strand reaction, we added 10 uL of in vitro transcription mixture using the HiScribe T7 High Yield RNA Synthesis Kit (E2040S, NEB), containing 1.5 uL of each NTP, 1.5 uL of T7 RNA Polymerase Mix, 1.5 uL of 10X T7 Reaction Buffer and 1 uL of RNaseOUT. The reaction was incubated at 37°C overnight (13-16 hours total). *Fragmentation reaction.* After the overnight incubation, 4 uL of ExoSAP-IT reagent (78201.1.ML, Applied Biosystems) was added and the mixture was incubated at 37°C for 15 minutes. Next, 2.67 uL of 10X RNA Fragmentation Buffer from the NEBNext Magnesium RNA Fragmentation Module (E6150S, NEB) was added, resulting in a total volume of 26.67 uL. The reaction was incubated in a preheated thermal cycler at 94°C for 2 minutes 50 seconds, followed by immediate transfer to ice and addition of 2.67 uL of 10X RNA Fragmentation Stop Solution. The reaction was then cleaned up with 41.1 uL of RNAClean XP beads (1.4X ratio), and resuspended in 5.5 uL of water.

#### Second reverse transcription reaction

To the 5.5 uL purified fragmentation reaction, we added 0.5 uL of 10 mM dNTP mix and 0.5 uL of 50 uM second RT primer for a total reaction volume of 6.5 uL. The reaction was incubated at 65°C for 5 minutes, then placed on ice for at least 1 minute. Next, we added 4 uL of RT reaction mix consisting of 2 uL of 5X First-strand Buffer, 1 uL of 100 mM DTT, 0.5 uL of RNaseOUT, and 0.5 uL of 200 U/uL SuperScript II Reverse Transcriptase (18064014, Invitrogen). The reaction was then incubated at 25°C for 10 minutes and 42°C for 1 hour.

#### Final PCR amplification

Using 5 uL of the second reverse transcription reaction, we prepared a 50 uL total volume PCR reaction containing 0.5 uM each of Nextera i5 and i7 indexed adapter primers, 1X Terra PCR Direct Buffer and 1 uL of 1.25 U/uL Terra PCR Direct Polymerase Mix (639270, Takara Bio). To clean up the reaction, 45 uL of RNAClean XP beads (0.9X ratio) was added, and after washes, libraries were eluted with 50 uL of water. A second round of bead purification was performed with the same bead ratio and the final purified library was eluted with 50 uL of water.

#### Library QC and sequencing

Final LSV-seq libraries were assessed using the Agilent High Sensitivity DNA Kit (5067-4626, Agilent) and quantified with the NEBNext Library Quant Kit for Illumina (E7630L, NEB). Libraries were then pooled and sequenced in 150-cycle single-end format on an Illumina NextSeq 550.

### Prioritization of targeted splicing events

#### Defining low-coverage, non-changing splicing events

To identify low-coverage splicing events which we could enrich with LSV-seq, we first ran the MAJIQ HET algorithm with parameters: {-- min-experiments 0.1} on GTEx samples across all 3 tissues, with the read coverage output enabled. This allowed us to identify target LSVs which were changing (at least 1 junction confidently changing) and non-changing (no junctions are confidently changing) across tissues, as well as to segregate high- and low-coverage LSVs based on a threshold of 50 mean reads of coverage. The voila modulize command was also run, which allowed us to extract the subset of LSVs located within cassette exons for the splicing code pipeline.

#### Prioritizing splicing events with CLIP-Seq data

Based on CLIP-Seq data in the K562 and HepG2 cell lines from ENCODE, we first identified putative “tissue-specific” RBPs. We filtered for the subset of available RBPs with at least 5 TPM mean expression in at least 1 of the 3 GTEx tissues of interest (brain cerebellum, heart atrial appendage, and liver) and exhibiting at a 2-fold change in expression in at least 1 pairwise tissue comparison. We then identified the top 50 RBPs ranked by their maximal log2 pairwise expression change across tissues, resulting in an initial shortlist of tissue-specific RNA-binding proteins.

Next, we discovered which of these top tissue-specific RBPs specifically regulate tissue-specific target LSVs across our 3 tissues of interest. For every high-coverage LSV we identified from our MAJIQ HET analysis we defined cis regulatory regions that might be bound by RBPs. This was defined as a combination of regions including every detected junction from 300 base pairs downstream to 50 base pairs upstream, and the 5’ boundary of the target exon itself from 50 base pairs downstream to 300 base pairs upstream. We intersected the called CLIP-Seq peaks for each tissue-specific RBP with the cis sequence regions of tissue-specific LSVs. Tissue-specific RBPs which putatively regulated tissue-specific splicing were defined as RBPs which bound changing LSVs more frequently than to non-changing LSVs, as evaluated by binomial tests for statistical significance.

Having defined tissue-specific splicing regulatory RBPs based on the above procedure, we were then able to define a candidate set of events to interrogate with LSV-seq. Specifically, this set consisted of the low-coverage LSVs that we failed to call confident changes for. The final candidate set was further required to overlap with CLIP-Seq peaks for at least 2, 4, or 6 such RBPs, depending on the stringency of our predictions.

#### Prioritizing splicing events with splicing code model

To prioritize splicing events using our deep learning splicing code model, we used our model to generate predictions for changes in mean PSI values for cassette exons detected in the MAJIQ HET analysis but suffered lack of coverage as described above.By tuning the threshold of predicted change between any pair of tissues to either 0.1, 0.15, or 0.2 PSI, we were able to vary the stringency of this pipeline.

### LSV-seq primer design

#### Obtaining candidate primer sequences from RNA-seq

To retrieve regions adjacent to known target LSVs (3’ splice sites), we ran the MAJIQ build and heterogen commands on the desired set of RNA-seq bam files, followed by the VOILA modulize command with the parameters{--keep-constitutive --decomplexify-psi-threshold 0 --show-all --output-mpe}. From the output file (’mpe_primerable_regions.tsv’), we used the columns for ‘Reference Exon Constant Region’, ‘Constitutive Regions’, ‘LSV ID’, ‘strand’, and ‘chromosome’ to create a BED file matching the LSV names to their genomic coordinates to be extracted. Prior to the next step, we obtained a list of repeats found in the hg38 genome from the RepeatMasker website^48^ and reformatted it as a BED file. We then used the bedtools^49^ subtract command to remove regions corresponding to the RepeatMasker defined repeat regions.

#### Generation of candidate primers and feature extraction

Primers were designed by running the primerGen script, which was run with default flags, to output primers with specific GC content (10-90%), melting temperature calculated based on RNA-DNA hybridization^50^ (50 to 85°C), length (15 to 40 bases), and lacking prohibited sequences (AAAAA, TTTTT, CCCCC, GGGGG). To obtain the full model specific features, BLAST^19^ was run against the human transcriptome with parameters {-gapopen 2 -gapextend 2 -reward 1 -penalty −2} and the NUPACK nucleic acid modeling package^20^ was run assuming reverse transcription reaction chemical and temperature conditions. For both the lite and full model, the featureExtract script was run to extract sequence-based features and append the alignment-based features from the previous step and experiment-specific features from externally provided files. The sequence-based features include encodings of positional nucleotide motifs including one-hot encodings of positional mononucleotides (e.g. presence of G at position 1 or A at position 2), one-hot encodings of positional dinucleotides (e.g. presence of TA at position 3 or presence of GG at position 5), and cumulative proportion of mononucleotides up to the i^th^ position (e.g. proportion of As within the first 5 positions). The resulting data matrix, consisting of rows of candidate primers and columns of features, was saved as a pickled file.

#### Prediction of primer performance

The modelPredict script was used to predict the performance of primers based on the saved data matrix of features. For the specified model version, either lite or full, predictions are outputted based on the saved yield/amplification model, denoted as a, and the specificity model, denoted as s. These predictions are combined into a single score using the following formula: (λ^a^)*s. The λ parameter controls the relative tradeoff between amplification and specificity, where higher λ increases the relative weight of the amplification prediction relative to the specificity prediction. In practice, λ was fixed to 1.2 in all the work presented here as it seemed to give a good tradeoff as reflected in the empirical results. With λ set, the primer with the maximal combined prediction score for each targeted region was outputted as the best predicted primer.

### Analysis of RNA-seq and LSV-seq data

#### Visualization and quantification of splicing events

For all analyzed LSV-seq and RNA-seq datasets, reads were first trimmed with BBDuk^51^ using the parameters {ref=adapters ktrim=r k=23 mink=11 hdist=1 tpe tbo qtrim=r trimq=15 qin=auto minlength=30}.

For RNA-seq analysis, we aligned reads to the hg38 genome using STAR^52^ with the parameters {-alignSJoverhangMin 8 --alignEndsType Local --outFilterMultimapNmax 1}. We then ran MAJIQ build across all samples in a given analysis with the parameters {--minreads 2 --min-denovo 2 --minpos 1 --target-lsvs}. For the Jurkat T-cell libraries, we ran MAJIQ deltapsi or psi with the parameters {--minpos 1 –minreads 2}, followed by visualization in the VOILA web browser viewer. For calculation of differential gene expression, summed TPMs for each gene were derived from transcript TPMs based on Kallisto^53^ quantifications and the mean was taken across all samples per condition. Subsequently, the log fold-change between the mean gene TPMs per condition was calculated.

For LSV-seq analysis, the umi-tools *extract* command^54^ was used to first extract the 10-nucleotide UMI from each read using the parameters {--bc-pattern=NNNNNNNNNN}, followed by STAR alignment with the same parameters described for RNA-seq. Then, umicollapse^55^ with parameter {--two-pass} was used to mark UMI read duplicate groups. Because the initial 5’ end primer sequence is also expected to be shared between identical reads, we appended the first 13 nucleotides beyond the UMI to the marked UMI sequences. Final deduplicated reads were selected using custom scripts which first filtered for reads which were at least 30 nucleotides in length, then selected one of the maximal length reads at random for each marked UMI group (consisting of both the marked UMI group itself and the downstream primer barcode). This resulted in the deduplicated BAM files we used in splicing visualization and downstream quantification.

To visualize splicing events, we used an adaptation of the MAJIQ deltapsi algorithm^12^, running MAJIQ build, and MAJIQ psi or deltapsi with the same parameters described for RNA-seq. We visualized the resulting splicing events in the VOILA web browser viewer. For RNA-seq, the classical MAJIQ algorithm integrates a Bayesian approach in order to remove potential read stacks which might correspond to PCR duplicates. However, because UMI deduplication is performed upstream of MAJIQ in LSV-seq, and because the expected read distribution for LSV-seq greatly differs from that expected for RNA-seq, we used a distinct version of the MAJIQ algorithm which takes the raw splice ratio per junction with the Bayesian modeling disabled. To quantify splicing, we ran the build command with the same corresponding parameters {--minreads 2 --mindenovo 2 --minpos 1} and output calculated splice ratios with the psi-coverage using parameters {--target-lsvs --stack-pvalue-threshold 0 --minbins 1 --minreads 2}. To quantify changes in expression, we took the total read count per LSV output from MAJIQ, normalized by total number of reads spanning targeted splicing events, as a proxy for overall gene abundance and calculated the log fold-change between LSV level read counts across conditions of interest.

When downsampling analyses were required for both RNA-seq and LSV-seq, we subsampled the trimmed read files to varying read depths prior to alignment and other downstream analyses. To account for the fact that RNA-seq was paired-end data, while LSV-seq was single-end, we filtered for RNA-seq reads with the “READ1” SAM flag set.

#### Analysis of on- and off-target reads per primer

To identify reads mapping on- and off-target, we created a BED file containing all of the on-target primer “extended” regions, defined as the region between the 3’ end of the primer, excluding the length of the primer itself, and the known 3’ splice site. We used this BED file to perform the bedtools intersect command on both RNA-seq and LSV-seq mapped deduplicated BAM files, retrieving the number of reads per on-target region. For each on-target region, fold enrichment over RNA-seq for each LSV-seq library was calculated by dividing the number of library-size normalized reads in LSV-seq by the average number of library-size normalized reads in RNA-seq replicates. To calculate on and off-target percentage, we used the bedtools intersect command to calculate the number of individual reads overlapping targeted regions, and divided by the total number of unique reads in the library.

### Training of Optimal Prime prediction models

#### Processing of output variables

To map each read to the original primers that they most likely originated from, the “blast_process” script was created. For each library, we first converted the BAM files mapped by STAR into SAM and FASTA files. The FASTA file was used to generate a BLAST database, which was aligned against the sequences of all the primers in the pool with parameters {-gapopen 2 -gapextend 2 -reward 2 -penalty 3 -task blastn-short}. For each sequenced library read, the primer that aligned within 50 base pairs of the 5’ read end with the highest alignment score was inferred to be the primer of origin. We removed unextended primer reads which were no more than 5 nucleotides longer than the primer itself.

While the original STAR alignment for each read was highly efficient at identifying the correct genomic positions to map, it often gave inaccurate information about insertions, deletions, or mutations near the 5’ read ends. To generate a more accurate representation of how the 5’ read ends mapped to their origin primers, we extracted the sequence corresponding to the mapped genomic coordinates of the read end. We then used the pairwise2.align.globalms method from the *biopython* package to align this sequence to the read’s origin primer sequence with highly permissive parameters {match=2, mismatch=-1, open=-0.5, extend=-0.2}. The top scoring alignment was taken as the most likely alignment of the actual genomic location of each read to its origin primer. The longest continuous sequence uninterrupted by insertions, deletions or mutations starting from the 3’ end of the primer, divided by the length of the primer itself is what we termed the “fraction of primer binding” (FPB).

Separately, we inferred whether a read was on- or off-target based on if the read overlapped with the extended primer region, corresponding to the region between the targeted 3’ splice site and the 5’ end of the primer, which includes the length of the primer itself. This allowed us to derive a specificity metric which we termed “on-target fraction” (OTF), corresponding to the number of on-target reads divided by the total number of reads for each primer. We also created a separate yield/amplification metric termed “log total amplification", corresponding to the logarithm of the total number of reads for each primer, first divided by the pool size and then divided by the library size. To aggregate multiple measurements of the same primer pool and tissue condition, we took the average of the specificity and amplification metrics. To account for redundant primers, especially when the same primer pool was used in a different tissue condition, we collapsed the feature vectors for these primers which were identical by sequence into a single mean vector.

#### Regression model for primer selection

Using the feature matrix we extracted during primer generation, and the output variables we processed, we trained CatBoost gradient boosting decision tree regression models, after iterating through different models and formulations of output variables. For the final specificity output variable, which was defined as the mean of FPB and OTF, we filtered for data points which had at least 12 total detected reads, as poorly amplified primers were subject to variance in measurement. In contrast, we did not use filtering for the final amplification output variable, defined as the log total amplification, because poorly amplified primers improved model performance. For each model, we performed 5-fold cross-validation twice and interpreted models using the SHAP TreeExplainer package^22^.

We also created two deep learning model architectures for predicting primer specificity and amplification. The transformer model, based on DNABERT^56^, was fine-tuned on mRNA transcripts and modified for regression. We used 6-mer sequences to train the transformer model. We also created a convolutional neural network (CNN) model, which utilized several convolution layers followed by a long short-term memory (LSTM) layer. Non-sequence-based features, such as expression, bitscore, and Tm, were concatenated before the second-to-last layer for both models. Both models were trained on the same primer data and underwent 5-fold cross-validation twice on identical splits as for the boosted decision tree models.

## Code Availability

All code for the analyses and processing pipelines and the link to the webtool will be made available upon publication.

## Acknowledgements

We thank the members of the Choi and Barash laboratories for helpful discussions. We want to specifically thank Farica Zhuang for assisting in development of the OP transformer model, Di Wu for providing splicing code predictions for event prioritization, and Joseph Aicher for implementation of the specific variant of the MAJIQv3 algorithm we used for LSV-seq analysis. We also thank Drs. Hagen Tilgner, Sydney Shaffer and Brian Gregory for valuable feedback. P.S.C. was supported by funding from the NCI (grant no. R00CA208028), the NIH Director’s New Innovator Award (grant no. DP2GM146251) and faculty startup funds from the Children’s Hospital of Philadelphia (CHOP). Y.B. was supported by funding from the NLM (grant no. R01-LM-013437) and NIGMS (grant no. GM128096).

