## Supplemental Figures for "Machine learning-optimized targeted detection of alternative splicing"

#### Figure S1

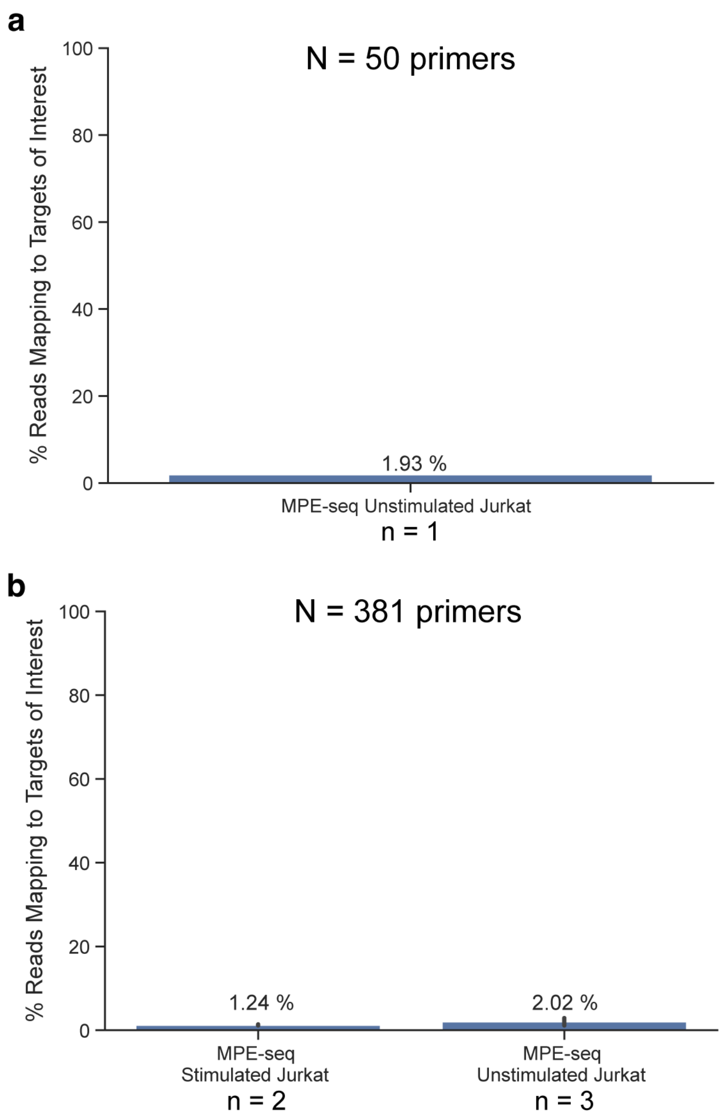


**Fig. S1 | Library preparation experiments in Jurkat T-cells using the original MPE-seq protocol.**

**a,** A 50-primer pool was used to target splicing events in a single unstimulated Jurkat T-cell sample. The resulting on-target percentage is shown. **b,** A 381-primer pool was used to target splicing events in both unstimulated (n=3) and stimulated (n=2) Jurkat T-cell sample replicates. The resulting mean on-target percentage per condition is shown.

**Figure S2**

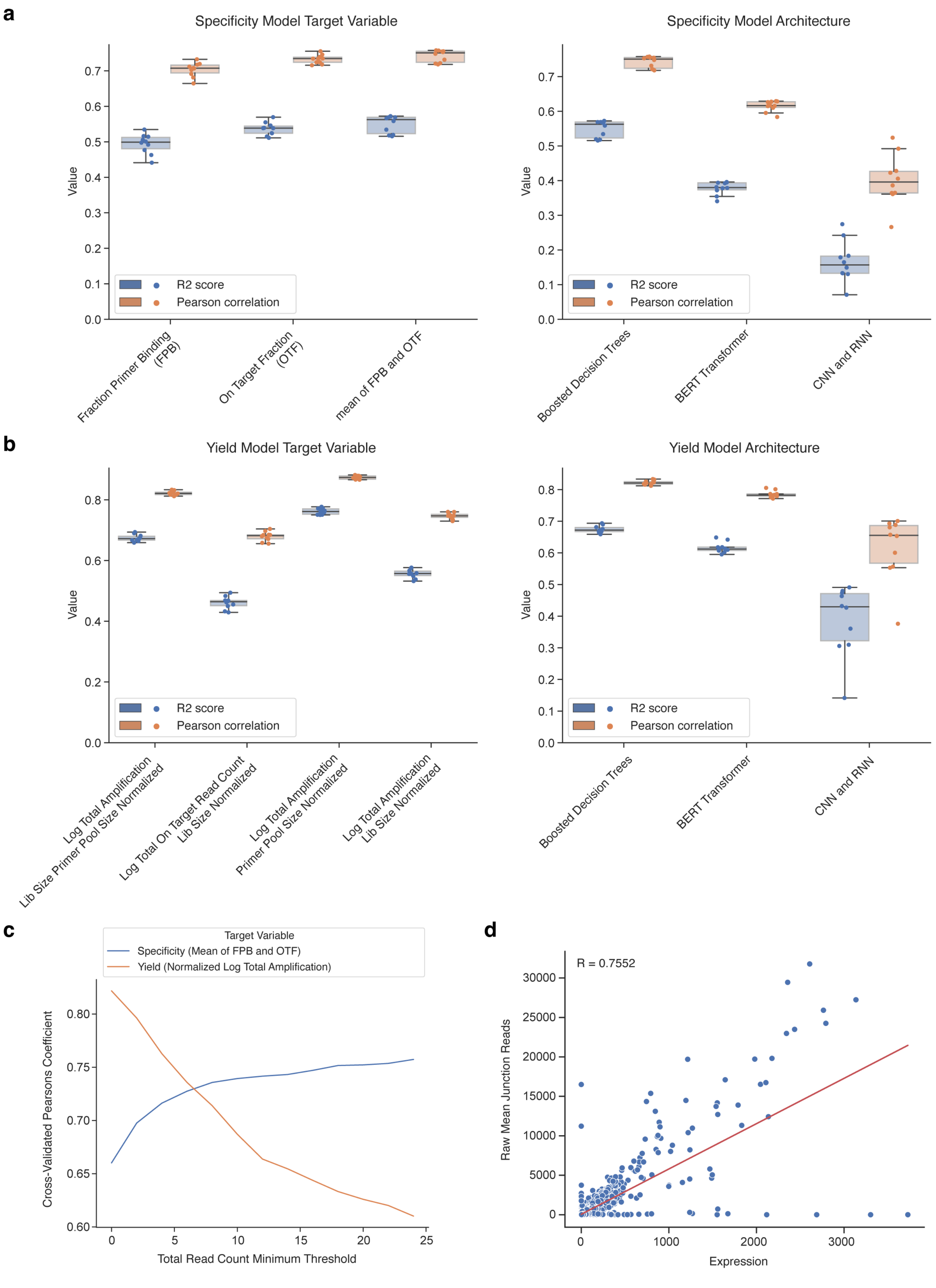

a
Specificity Model Target Variable
Value
R2 score
Pearson correlation
Fraction Primer Binding (FPB)
On Target Fraction (OTF)
mean of FPB and OTF
Specificity Model Architecture
Boosted Decision Trees
BERT Transformer
CNN and RNN
b
Yield Model Target Variable
Log Total Amplification Lib Size Primer Pool Size Normalized
Log Total On Target Read Count Lib Size Normalized
Log Total Amplification Primer Pool Size Normalized
Log Total Amplification Lib Size Normalized
Yield Model Architecture
c
Target Variable
Specificity (Mean of FPB and OTF)
Yield (Normalized Log Total Amplification)
Cross-Validated Pearsons Coefficient
Total Read Count Minimum Threshold
d
R = 0.7552
Raw Mean Junction Reads
Expression

**Fig. S2 | Training considerations during development of Optimal Prime models.**

**a,** For the specificity models, the effects of different tested conditions during model training on prediction accuracy, across 5 cross-validated splits performed independently twice, shown as box plots for both R2 score and Pearson correlation metrics. We considered multiple different formulations of the predicted variable including FPB, OTF, and the mean of FPB and OTF. We also considered different model architectures including boosted decision trees, BERT transformer, and CNN and RNN. **b,** For the yield models, the effects of different tested conditions during model training on prediction accuracy, across 5 cross-validated splits performed independently twice, shown as box plots for both R2 score and Pearson correlation metrics. We considered multiple different formulations of the predicted variable including various different normalization methods for the logarithmic total amplification. We also considered different model architectures including boosted decision trees, BERT transformer, and CNN and RNN. **c,** For both the specificity (blue line) and yield (orange line) models, line graphs depict the effects of filtering our datasets by minimum thresholds of total read coverage. Because the read coverage filter only improved performance for the specificity models, we omitted it for the final yield models. **d,** Scatter plot depicting linear relationship modeled between expression and raw junction read counts. For a subset of LSVs which were not detected and had an effective junction read coverage of zero, we had to fill in missing junction count values to complete the dataset prior to training. To accomplish this, we modeled junction count values as a linear regression on overall gene expression.

### Figure S3

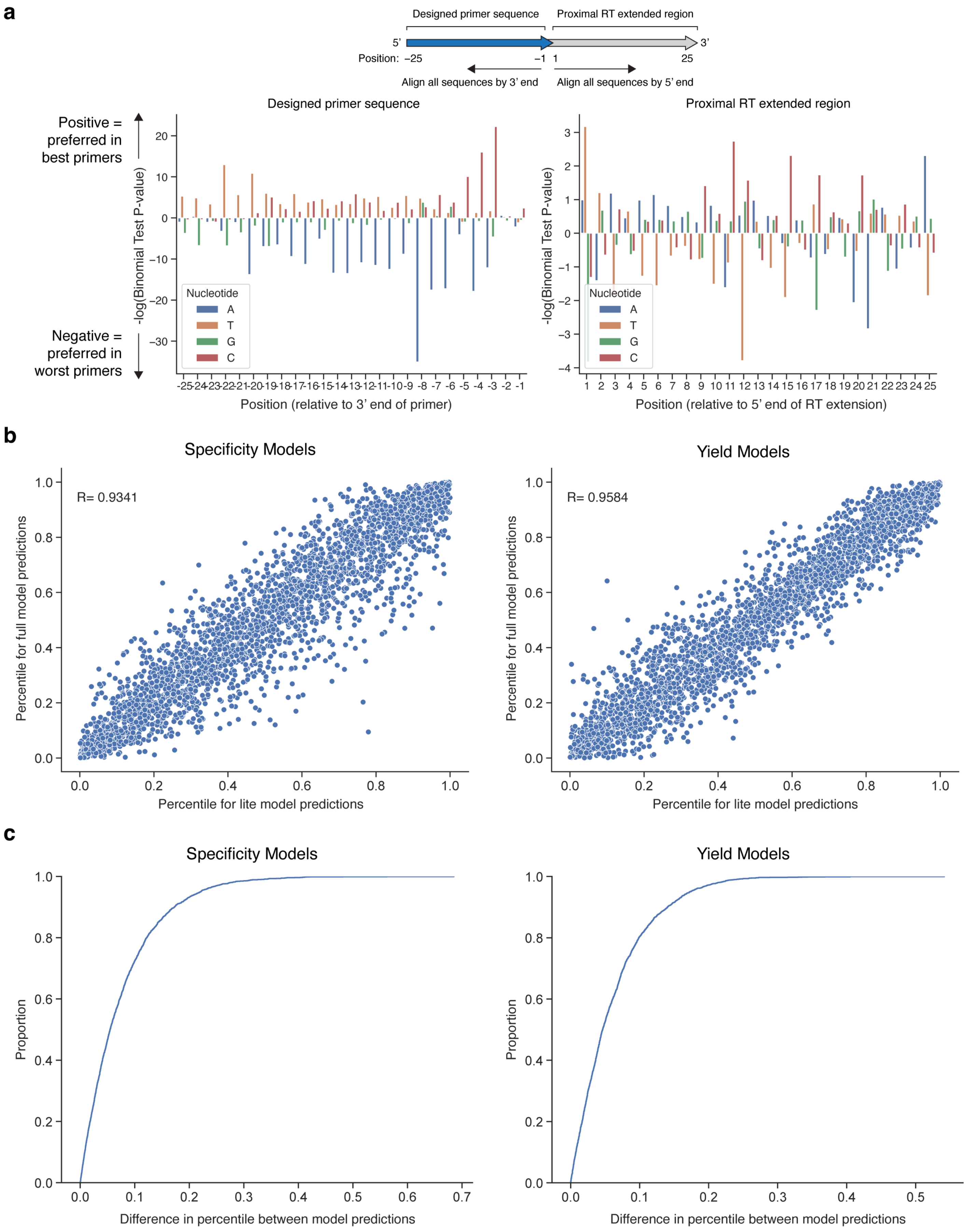
a
Designed primer sequence
Proximal RT extended region
5'
3'
Position: –25
–1
1
25
Align all sequences by 3' end
Align all sequences by 5' end
Designed primer sequence
Proximal RT extended region
Positive = preferred in best primers
Negative = preferred in worst primers
-log(Binomial Test P-value)
Nucleotide
A
T
G
C
Position (relative to 3' end of primer)
Position (relative to 5' end of RT extension)
b
Specificity Models
Yield Models
R= 0.9341
R= 0.9584
Percentile for full model predictions
Percentile for lite model predictions
c
Specificity Models
Yield Models
Proportion
Difference in percentile between model predictions

**Fig. S3 | Additional evaluation and interpretation of Optimal Prime models.**

**a,** Similar to Figure 3d for the specificity metric, bar graphs depicting the relative enrichment of specific nucleotides at the first 25 positions in both directions from the 3' end of the primer, for either the best or lowest performance primer groups, as evaluated by the yield metric (Normalized Log Total Amplification). For each position and nucleotide combination, a binomial test was conducted to determine the likelihood of the observed proportion in the top quintile of primers, compared to the proportion in the bottom quintile of primers. **b,** For specificity and yield models, scatter plots depicting the relative consistency of the lite and full model versions for ranking primers by percentile. Models were each retrained while holding out 20% of the dataset selected at random as the test set. Then, model predictions for the lite and full models were independently obtained for the test set, the predicted values were converted into percentile based rankings, and the resulting percentiles for each data point were plotted for each model. **c,** For specificity and yield models, the cumulative distribution function depicting the distribution of differences in percentile ranking between the lite and full model versions, based on the held out test set as in b.

### Figure S4

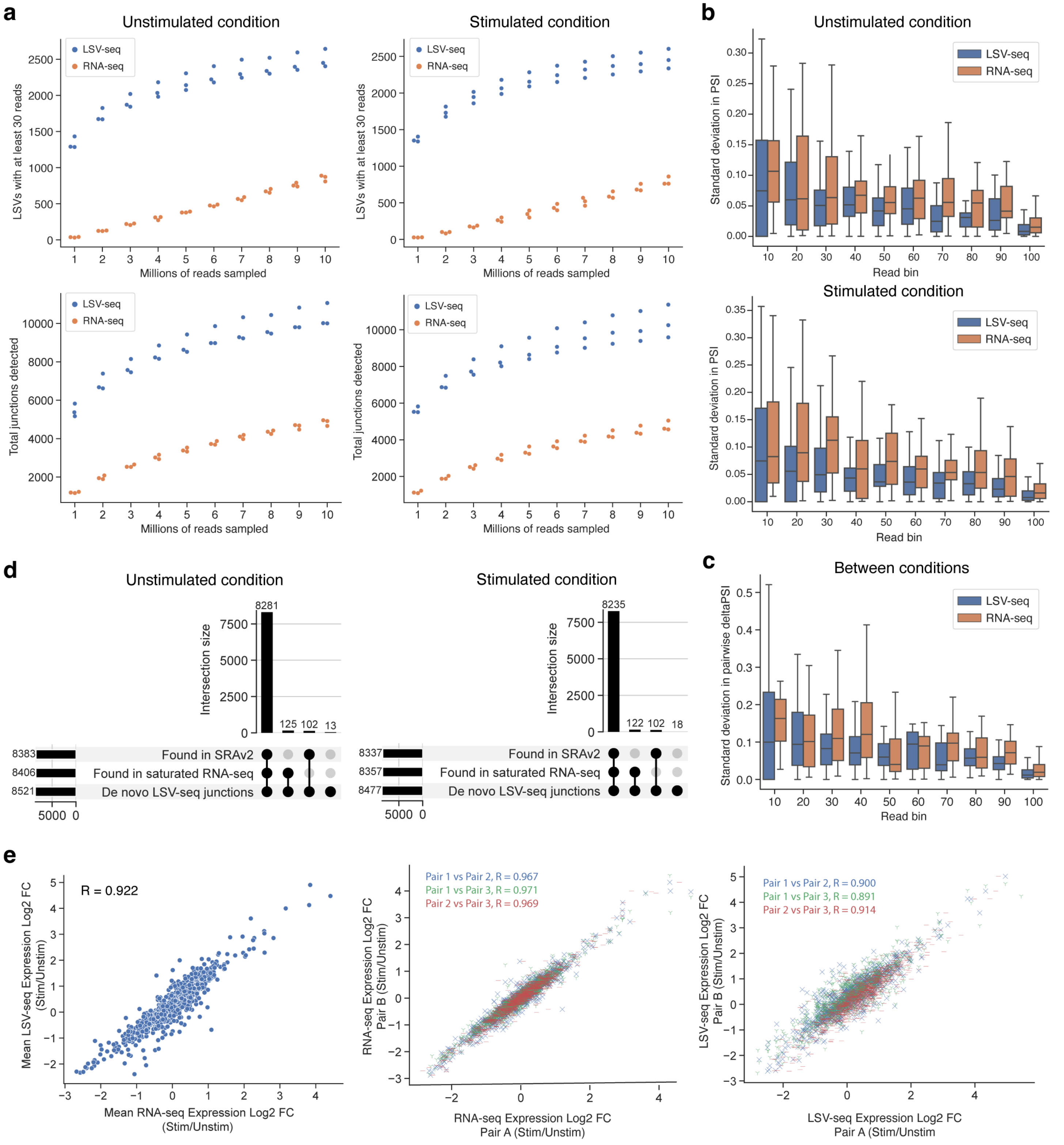


**Fig. S4 | Additional details of LSV-seq benchmark analyses in Jurkat T-cells.**

**a,** For unstimulated and stimulated Jurkat T-cell LSV-seq and RNA-seq datasets, the effects of different levels of downsampling shown as dot plots for two different metrics: the number of LSVs with at least 30 reads (top) and the total number of detected junctions (bottom). For the remaining downsampling analyses throughout the text, we focused on datasets downsampled to a depth of 3 million reads. **b,** Box plots showing the variance in

PSI value measurements across different read coverage bins for unstimulated and stimulated Jurkat T-cell LSV-seq and RNA-seq datasets. We calculated the standard deviation in PSI values for every LSV across 3 biological replicates. Because variance in measurements is expected to correlate with mean read coverage, we then binned each LSV based on mean read coverage. **c,** Box plot showing the variance in pairwise deltaPSI value measurements across different read coverage bins. For each matched unstimulated and stimulated Jurkat T-cell sample pair in the LSV-seq and RNA-seq datasets, we calculated the measured difference in PSI values for every LSV, which we termed the pairwise deltaPSI. Then, we calculated the standard deviation in pairwise deltaPSI for LSVs which were detected in all 3 Jurkat T-cell sample pairs and binned each LSV based on mean read coverage. **d,** Upset plots showing that de novo LSV-seq junctions are present in higher coverage datasets. For the downsampled unstimulated and stimulated Jurkat T-cell LSV-seq and RNA-seq datasets, we isolated LSV-seq junctions which were only found in LSV-seq but not RNA-seq (“De novo LSV-seq junctions”). Then, we asked if these junctions could be found in any of the saturated RNA-seq sample datasets (“Found in saturated RNA-seq”) or found in a collection of all junctions found in the SRA, accessed via the Snaptron database[43] (“Found in SRAv2”). **e,** Scatter plots comparing the stimulated - unstimulated log2 expression difference for the same genes in either full-depth LSV-seq or RNA-seq datasets. Pearson correlation coefficient is given. The first plot shows the mean LSV-seq versus RNA-seq expression difference, while the subsequent plots show the internal consistency of each pairwise unstimulated/stimulated differential expression, for both RNA-seq and LSV-seq.

### Figure S5

**a**

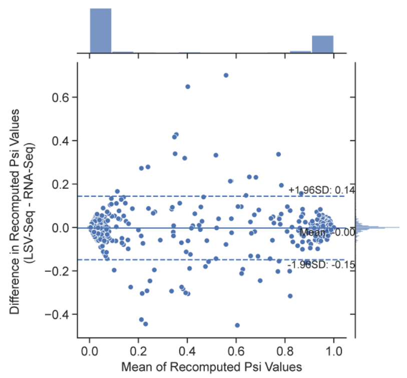


**b**

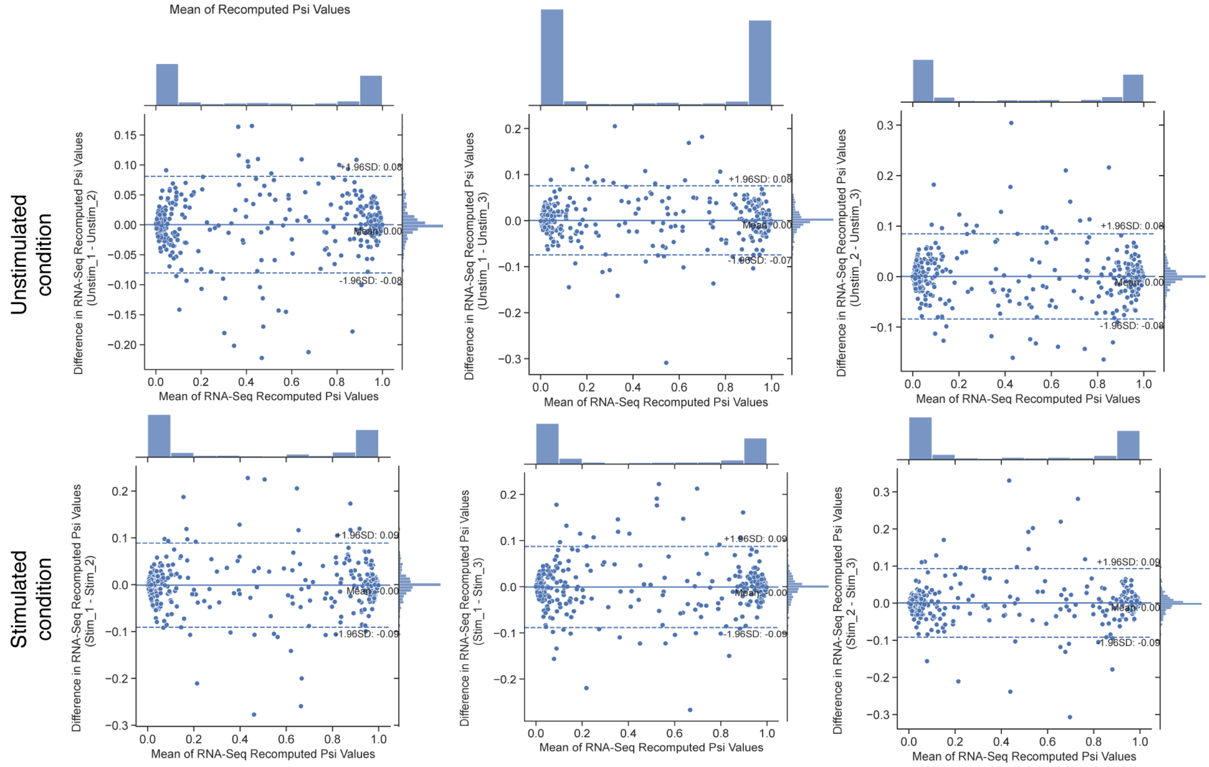


**c**

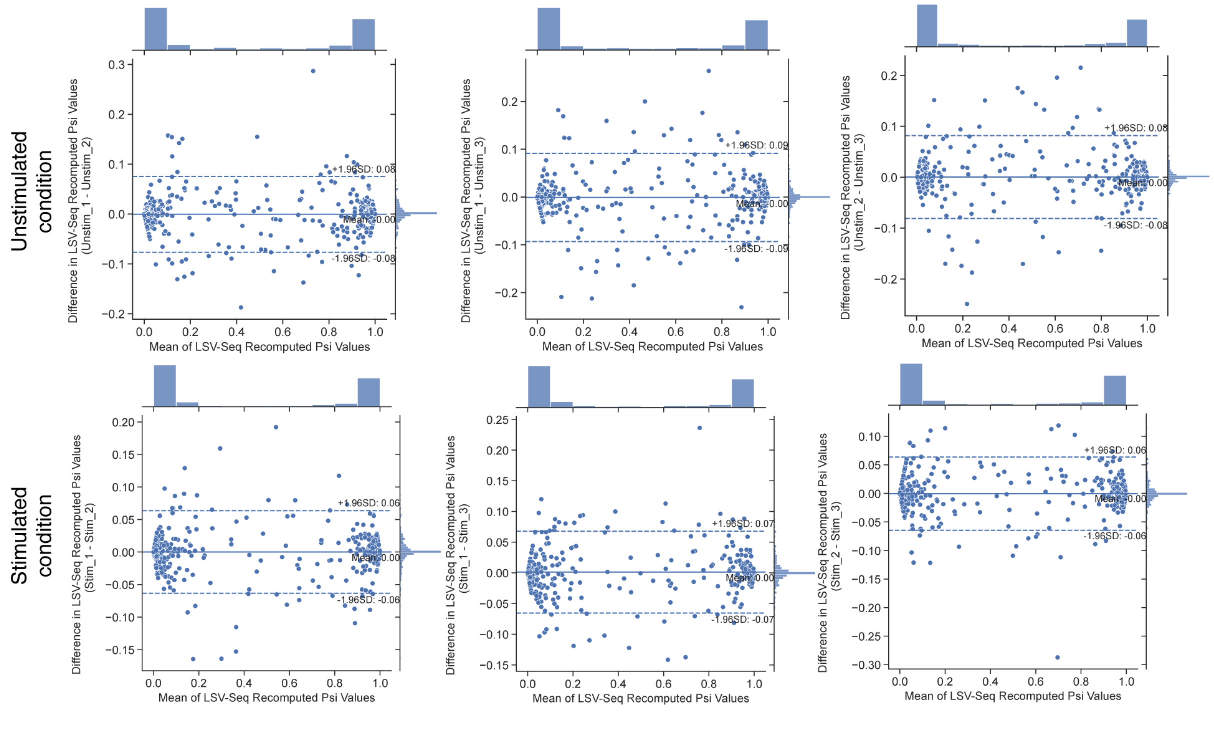

**Fig. S5 | Additional analyses of inter- and intra-method PSI value consistency.**

**a**, Bland-Altman plot comparing the inter-method consistency of LSV-seq and RNA-seq mean PSI value estimates, using the same data as Figure 4e. **b**, Bland-Altman plots comparing the intra-method consistency of individual RNA-seq sample PSI value estimates. **c**, Bland-Altman plots comparing the intra-method consistency of individual LSV-seq sample PSI value estimates.

### Figure S6

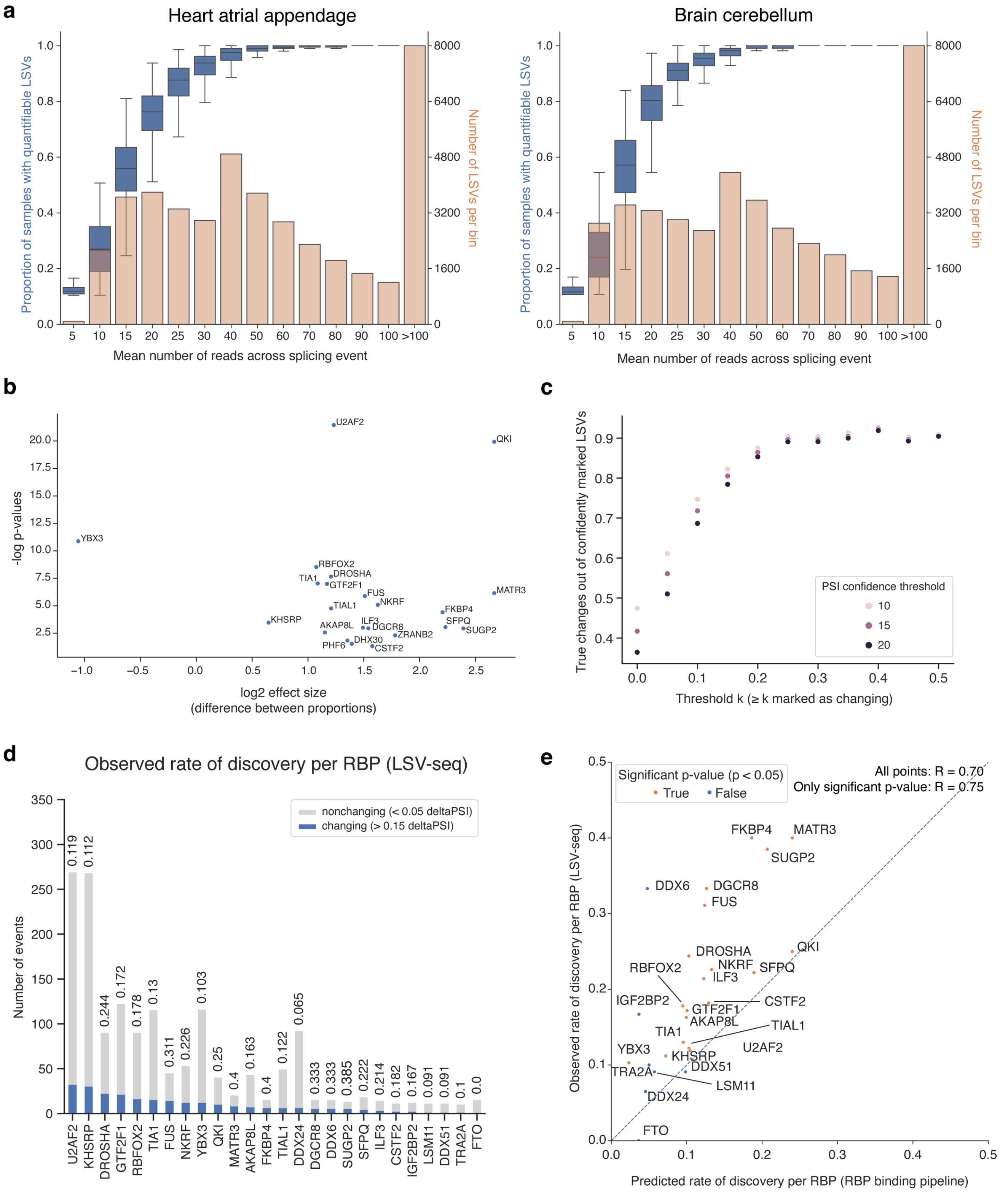
a
Heart atrial appendage
Brain cerebellum
Proportion of samples with quantifiable LSVs
Number of LSVs per bin
Mean number of reads across splicing event
b
-log p-values
log2 effect size
(difference between proportions)
U2AF2
QKI
YBX3
RBFOX2
DROSHA
TIA1
GTF2F1
FUS
NKRF
TIAL1
MATR3
FKBP4
SFPQ
SUGP2
KHSRP
AKAP8L
ILF3
DGCR8
ZRANB2
DHX30
PHF6
CSTF2
c
True changes out of confidently marked LSVs
Threshold k (≥ k marked as changing)
PSI confidence threshold
10
15
20
d
Observed rate of discovery per RBP (LSV-seq)
Number of events
nonchanging (< 0.05 deltaPSI)
changing (> 0.15 deltaPSI)
e
Significant p-value (p < 0.05)
True
False
All points: R = 0.70
Only significant p-value: R = 0.75
Observed rate of discovery per RBP (LSV-seq)
Predicted rate of discovery per RBP (RBP binding pipeline)

**Fig. S6 | Additional details of GTEx datasets and candidate prioritization pipeline.**

**a,** Similar to Figure 5b for liver, histograms in orange depicting distributions of mean read coverage across all detectable LSVs for heart atrial appendage and brain cerebellum. Box plots in blue are further overlaid to demonstrate the dropoff in proportion of quantifiable LSVs at lower read coverage levels. **b,** Volcano plot depicting the association between log2 effect size of tissue-specific RBPs, defined as the difference in proportion between known high-coverage tissue-specific and tissue-nonspecific LSVs, and their associated -log p-values, which assesses the statistical significance of the difference. **c,** Dot plot depicting the change in sensitivity across thresholds of predicted deltaPSI changes, of varying stringency, for the validation set of known high-coverage, tissue-specific LSVs. **d,** Stacked bar plots listing the discovery rates for individual RBPs, ordered by the absolute number of changing events which were returned by LSV-seq. **e,** Scatter plot depicting the correlation between predicted and observed LSV-seq discovery rates for individual RBPs. RBPs are colored based on whether or not they passed a significance p-value threshold in the RBP binding pipeline. Pearson correlation coefficient is given for both all RBPs and only the subset of RBPs with significant p-values.

Figure S7

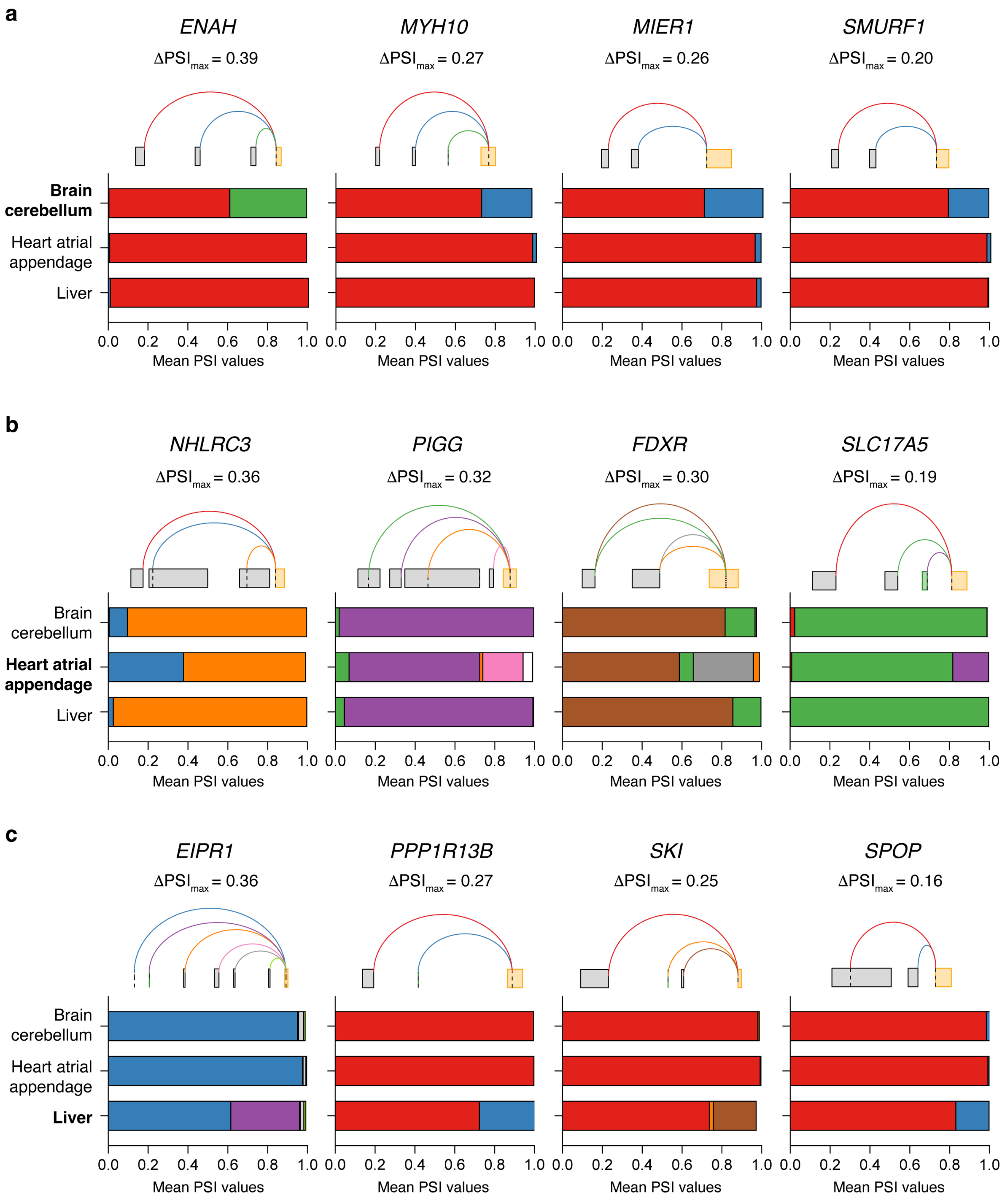
a
ENAH
ΔPSImax = 0.39
MYH10
ΔPSImax = 0.27
MIER1
ΔPSImax = 0.26
SMURF1
ΔPSImax = 0.20
Brain cerebellum
Heart atrial appendage
Liver
0.0 0.2 0.4 0.6 0.8 1.0
Mean PSI values
b
NHLRC3
ΔPSImax = 0.36
PIGG
ΔPSImax = 0.32
FDXR
ΔPSImax = 0.30
SLC17A5
ΔPSImax = 0.19
c
EIPR1
ΔPSImax = 0.36
PPP1R13B
ΔPSImax = 0.27
SKI
ΔPSImax = 0.25
SPOP
ΔPSImax = 0.16

**Fig. S7 | Additional tissue-specific splicing events recovered by LSV-seq.**

Examples of tissue-specific LSVs detected by LSV-seq with splicing unique to brain cerebellum (**a**), heart atrial appendage (**b**) or liver (**c**). Shown are stacked bar plots of junction PSI values and a Voila splice graph for each LSV, with colors of splice junctions corresponding to colors in PSI bar plots. DeltaPSImax refers to the maximum difference in PSI from among all splice junctions and pairwise tissue comparisons.
